# Preventing CpG island hypermethylation in oocytes safeguards mouse development

**DOI:** 10.1101/2024.05.29.595726

**Authors:** Yumiko K. Kawamura, Evgeniy A. Ozonov, Panagiotis Papasaikas, Takashi Kondo, Nhuong V. Nguyen, Michael B. Stadler, Sebastien A. Smallwood, Haruhiko Koseki, Antoine H.F.M Peters

**Author notes:** Shared second authorship (Evgeniy A. Ozonov, Panagiotis Papasaikas).

## Abstract

In mammalian somatic and male germline cells, genomes are extensively DNA methylated (DNAme). In oocytes, however, DNAme is largely limited to transcribed regions only. Regulatory CpG-island (CGI) sequences are also devoid of repressive DNAme in somatic and germ cells of both sexes. The mechanisms restricting *de novo* DNAme acquisition in developing oocytes, at CGIs and globally, and the relevance thereof for regulating zygotic gene expression and embryo development after fertilization are largely unknown. Here we show that the histone H3 lysine 36 dimethyl (H3K36me2) demethylases KDM2A and KDM2B prevent genome-wide accumulation of H3K36me2, thereby impeding global DNMT3A-catalyzed *de novo* DNAme, including at CGI gene promoters. By recruiting variant Polycomb Repressive Complex 1 (vPRC1), they further control H2A mono-ubiquitin deposition and vPRC1-dependent gene repression. Through genetic perturbations, we demonstrate that aberrant *Dnmt3a*-dependent DNAme established in *Kdm2a/Kdm2b* double mutant oocytes represses transcription from maternal loci in two-cell embryos. The lethality of *Kdm2a/Kdm2b* maternally deficient pre-implantation embryos is suppressed by *Dnmt3a* deficiency during oogenesis. Hence, KDM2A/KDM2B are essential for confining the oocyte DNA methylome, conferring competence for early embryonic development. Our research implies that the reprogramming capacity eminent to early embryos is insufficient to erase aberrant DNAme from maternal chromatin, and that early development is vulnerable to gene dosage haplo-insufficiency effects.

**HIGHLIGHTS:** Demethylation of H3K36me2 by KDM2A and KDM2B prevents aberrant de novo DNA methylation in mouse oocytes.

Sequence composition and H3K4me3 modulate the probability for aberrant H3K36me2 and DNA methylation at CpG islands.

Aberrant oocyte DNA methylation is not reprogrammed in early embryos and suppresses maternal gene transcription.

Aberrant oocyte DNA methylation causes embryonic lethality during pre-implantation development.

**GRAPHICAL SUMMARY:** 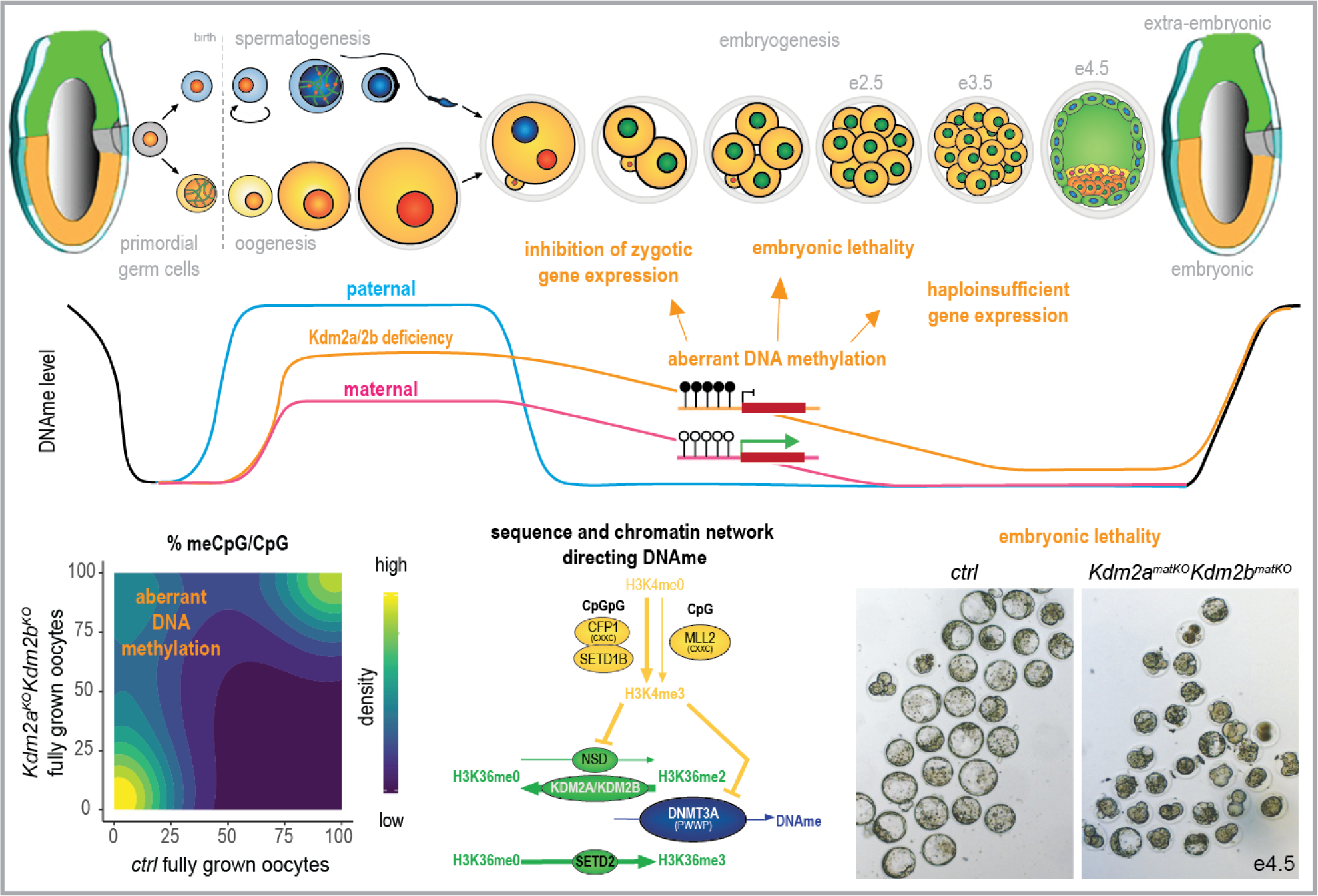

## INTRODUCTION

Shortly after fertilization, parental genomes undergo extensive reprogramming of germline specific epigenetic programs, including DNA methylation and chromatin, to support acquisition of totipotency and embryonic development. DNA methylation in mammals is generally found on cytosine residues within CpG dinucleotides (mCpG) throughout the genome, while CpGs located within CpG dinucleotide-dense regions, commonly referred to as CpG islands (CGIs), are generally unmethylated throughout the entire mammalian life cycle, including in sperm and oocytes. Many CGIs serve transcriptional regulatory functions at housekeeping and cell fate determining genes^1^. Through evolution, CpGs have become underrepresented in mammalian genomes due to deamination and incorrect repair of methylated cytosines in the germline^2^. To date, the mechanisms ensuring the unmethylated status of CGIs in the germline remain poorly understood. It is further unknown whether the germline-derived unmethylated state of CGIs in parental genomes is required for zygotic genome activation in early embryos and for supporting embryonic development, or whether (experimentally induced) DNAme at CGIs would undergo epigenetic reprogramming after fertilization.

Within the mammalian life cycle, genomes undergo two rounds of erasure and re-establishment of global DNAme patterns, largely excluding CGIs^3^. Following the specification of primordial germ cells, genomes first lose embryonically established DNAme patterns and then acquire oocyte and sperm specific patterns in a sexually dimorphic manner supporting germline specific cellular physiology. Next, following fertilization, both parental genomes lose most DNAme in a parent-of-origin specific manner^4^. After implantation, parental genomes become similarly and widely methylated during gastrulation. Despite extensive DNAme reprogramming during pre-implantation development, a few regions on maternal and paternal genomes escape erasure^5,6^. The transmission of DNAme at so-called imprinting control regions (ICRs) drives parent-of-origin specific mono-allelic repression which is vital to the developing embryo^3^.

Remarkably, global patterns and functions of DNAme in sperm and oocytes are very different. Male germ cells gain DNAme genome wide at over 90% of individual CpGs, comparable to levels in somatic cells. Such DNAme is essential for meiotic progression^7^ and maintenance of long-term spermatogenesis^8^. In contrast, growing oocytes acquire high levels of *de novo* DNAme exclusively in transcribed regions and only low levels in other regions, resulting in a global CpG methylation level below 40%^9,10,11^. Curiously, DNAme in oocytes does not majorly regulate gene expression nor is it required for oocyte development^12^. After fertilization, however, embryos lacking maternal DNAme arrest by day 10.5 of gestation, fairly late in development, due to genomic imprinting defects and/or impaired trophoblast formation^12,13^.

As in somatic cells, *de novo* DNAme acquisition in germ cells is controlled by histone methylation modifiers^14,15,16,17,18^. In growing oocytes, most DNAme catalyzed by the *de novo* DNA methyltransferase DNMT3A is directed by transcription-coupled H3K36me3, deposited by SETD2^19,20,9^. Moderate to low DNAme has been associated with H3K36me2 occupancy^11^. Inversely, DNMT3A catalysis is inhibited by H3K4me3 which is widely deposited in the oocyte genome by the SETD1B and MLL2 enzymes^21,22,23,24^. At selective regions including ICRs, however, H3K4me3 is removed by the KDM1B demethylase, enabling DNAme acquisition^22^. In mouse embryonic stem cells (ESCs), loss of KDM2B expression (also termed FBXL10, NDY1, JHDM1B and CXXC2) resulted in aberrant DNAme at certain CGIs controlled by Polycomb Repressive Complexes ^25^. The mechanism underlying such selective DNAme acquisition has, however, remained unknown^25^. Importantly, KDM2B localizes at almost all CGIs throughout the mouse genome via recognition of unmethylated CpGs by its CXXC domain^26^. The protein also contains a JmjC domain that was reported to demethylate H3K36me2 *in vitro*^27,28^. In ESCs, however, only limited activity was reported^29^. KDM2B further contains a PHD domain, an F-box domain and a leucine-rich repeat (LRR) that interacts with members of the variant Polycomb Repressive Complex 1.1 (vPRC1.1)^30,31,32^. vPRC1.1 deposits histone H2A mono-ubiquitin (H2AK119u1) at CGIs through the E3 ligases RING1 and RNF2, which are PRC1 core components, and is essential for transcriptional repression of target genes ^29,33^. Like KDM2B, the KDM2A paralog (FBXL11, JHDM1A, CXXC8) localizes at unmethylated CGIs^34^ and demethylates H3K36me2 *in vitro*^28^. KDM2A plays, however, only a minor gene regulatory function in ESCs, compared to KDM2B^29^.

We previously identified *Ring1* and *Rnf2* as critical transcriptional regulators and chromatin modifiers in oocytes, essential for defining embryonic competence^35^. Deficiency for *Pcgf1*, another component of vPRC1.1, indicated a role for this complex in defining H2AK119u1 and transcriptional states in oocytes^36^. Here, we study the role of *Kdm2b* and its paralog *Kdm2a* in regulating PRC1-mediated gene repression and *de novo* DNAme acquisition during oogenesis and the impact of their loss-of-function on embryogenesis^34,28^. We identify KDM2A and KDM2B as essential regulators of pre-implantation development, safeguarding the maternal genome against CpG hypermethylation throughout the genome including CGIs, thereby enabling proper zygotic gene expression and embryo viability.

## RESULTS

### *Kdm2a*/*Kdm2b* function in oocytes controls embryogenesis

RNA-sequencing experiments show that *Kdm2b* and other vPRC1 components are highly expressed in growing and fully grown germinal vesicle oocytes (GOs and FGOs) and in early embryos (Figure S1A). To study the function of KDM2B in oocytes and pre-implantation embryos, we conditionally altered its expression in GOs in two ways using the *Zp3*-promoter-driven CRE-recombinase expressed in primary GOs (Figure S1B). Firstly, removal of exons encoding the histone demethylase JmjC domain (*Kdm2b^fl-JmjC^*)^29^ fully abrogated *Kdm2b* expression (Figure S1C). We therefore refer to the *Kdm2b^fl-JmjC^* allele as a knock-out (*Kdm2b^KO^*) allele (Figure 1A). Secondly, we generated mice expressing a KDM2B protein lacking its CxxC domain (*Kdm2b^ΔCxxC^*), shown in ESCs to be unable to recruit vPRC1.1 to CGIs^31^ (Figures 1A and S1C). In both models, we did not observe changes in the number of ovulated oocytes nor in the progression of pre-implantation embryonic development after fertilization with wild-type (*wt*) sperm generating so called *Kdm2b^matKO^* or *Kdm2b^matΔCxxC^* embryos (Figures 1B and S1E).

**Figure 1:**
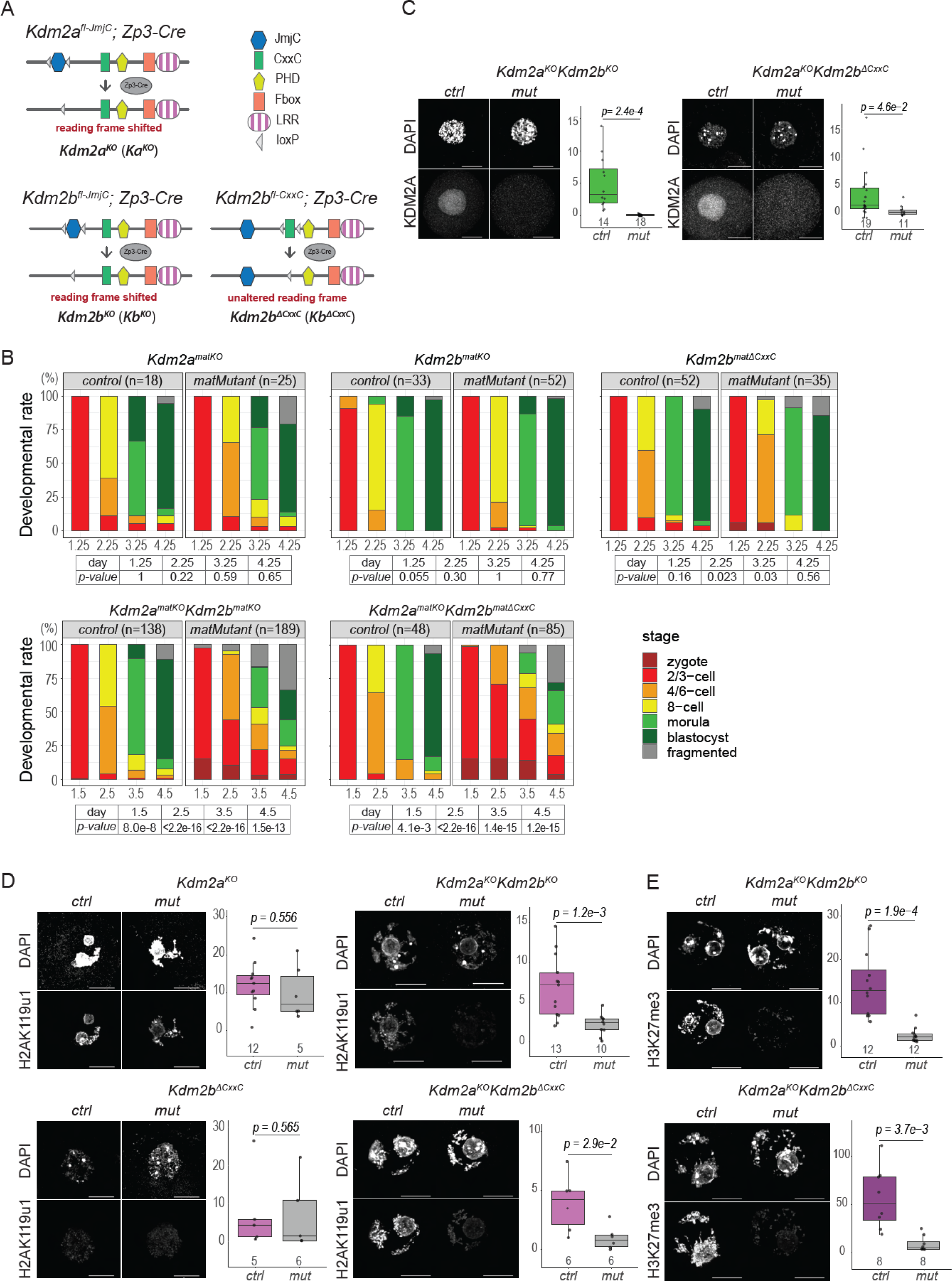
KDM2A and KDM2B function redundantly in oocytes to regulate embryonic development. **A.** Schematic overview of KDM2A and KDM2B proteins expressed in oocytes of *ctrl* and mutant conditions. Position of JmjC and CxxC domains flanked by loxP sites on the genome are indicated. Floxed alleles were deleted by CRE-recombinase, expressed from a *Zona pellucida 3*-cre (*Zp3*-cre) transgene initiated in primary GOs (see also Figure S1B) ^63^. Deletion of the JmjC domain encoding exons of *Kdm2a^fl-JmjC^* and *Kdm2b^fl-JmjC^* ^29^ results in a translational frame shift, decay of mRNA transcripts and greatly reduced expression. Deletion of the CGI-binding Zinc Finger Domain “CxxC” encoding exons of *Kdm2b^fl-CxxC^* ^31^ causes an in-frame excision. leading to expression of a slightly smaller KDM2B protein unable to be recruited to CpG islands (see also Figures S1C and S1D) ^31,29^. **B.** Developmental progression rates of pre-implantation embryos at embryonic day e1.25, e2.25, e3.25 and e4.25 (single mutants) or at embryonic day e1.5, e2.5, e3.5 and e4.5 (double/compound mutants), generated by fertilizing either *Kdm2a^KO^*, *Kdm2b^KO^*, *Kdm2b^ΔCxxC^* single mutant, *Kdm2a^KO^Kdm2b^KO^* (abbreviated as *Ka^KO^Kb^KO^*) double mutant, *Kdm2a^KO^Kdm2b^ΔCxxC^* (abbreviated as *Ka^KO^Kb^ΔCxxC^*) compound mutant, or respective *ctrl* oocytes with *wt* sperm and cultured over 4 days *in vitro*. Numbers of analyzed embryos are indicated. P-values according to Fisher’s exact test. **C.** Immunofluorescence staining and quantification of KDM2A protein expression in *Kdm2a^KO^Kdm2b^KO^*, *Kdm2a^KO^Kdm2b^ΔCxxC^* and *ctrl* GOs. The number of analyzed oocytes are indicated. P-values according to two-sided student’s *t*-test. **D.** Immunofluorescence staining and quantification of H2AK119u1 in *Kdm2a^KO^*, *Kdm2b^ΔCxxC^*, *Kdm2a^KO^Kdm2b^KO^*, *Kdm2a^KO^Kdm2b^ΔCxxC^* and respective *ctrl* FGOs. The number of analyzed oocytes are indicated. P-values according to two-sided student’s *t*-test. **E.** Immunofluorescence staining and quantification of H3K27me3 in *Kdm2a^KO^Kdm2b^KO^*, *Kdm2a^KO^Kdm2b^ΔCxxC^* and *ctrl* FGOs. Numbers of analyzed oocytes are indicated. P-values according to the two-sided student’s *t*-test.

To address developmental roles of the paralogous KDM2A, also relative to KDM2B function, we generated oocytes conditionally deficient for *Kdm2a* transcript and protein expression, alone or in combination with either *Kdm2b* mutation (Figures 1A, 1C and S1D). Whereas development of *Kdm2a^KO^* oocytes and resulting *Kdm2a^matKO^* embryos were not affected, double *Kdm2a^KO^Kdm2b^KO^* deficiency in oocytes severely impaired developmental progression of *Kdm2a^matKO^Kdm2b^matKO^* embryos towards the blastocyst stage, even though ovulation rates were normal (Figures 1B and S1E). Such embryonic impairment was phenocopied by embryos maternally compound mutant for *Kdm2a^KO^* and *Kdm2b^ΔCxxC^* (Figures 1B and S1E). Therefore, we conclude that KDM2A and KDM2B serve, additively or redundantly, essential functions during oogenesis to support pre-implantation development.

### KDM2A/KDM2B regulate PRC1-mediated gene repression

To assess the role of KDM2A/KDM2B in Polycomb-mediated functions, we first quantified the levels of PRC1-catalyzed H2AK119 mono-ubiquitination (H2AK119u1) in *ctrl* and mutant FGOs by immunofluorescence (IF) analyses. Whereas H2AK119u1 levels were not altered in FGOs singly deficient for either *Kdm2a* or *Kdm2b^ΔCxxC^*, they were massively reduced in *Kdm2a^KO^Kdm2b^KO^* and *Kdm2a^KO^Kdm2b^ΔCxxC^* FGOs (Figure 1D). These results argue that either KDM2A or KDM2B is sufficient for recruiting vPRC1 to chromatin, likely via their CxxC domains, to enable H2AK119u1 deposition. Moreover, PRC2-catalyzed H3K27me3 levels were greatly reduced in both types of double mutant FGOs (Figure 1E), indicating that vPRC1 functions upstream of PRC2 in GOs, as reported for ESCs^31^.

To understand the transcriptional regulatory function of KDM2A/KDM2B, we profiled RNA transcriptomes in single FGOs, from single and double mutants (Figures S2A and S2B). Compared to *ctrl* oocytes, over 1400 genes were upregulated in *Kdm2a^KO^Kdm2b^KO^* and *Kdm2a^KO^Kdm2b^ΔCxxC^* FGOs, while ∼550 genes were downregulated (Figures 2A, 2B, 2H and S3B). Most promoters of the 974 genes commonly upregulated are marked by H2AK119u1 in *ctrl* oocytes^36^ (Figure 2C, Tables S1 and S2). In addition, 71% of genes upregulated in *Kdm2a^KO^Kdm2b^KO^* FGOs were also upregulated in FGOs deficient for *Ring1* and *Rnf2*, two core components of all PRC1 complexes, acting redundantly during oogenesis^35^ (Figures 2D, 2E, 2H, Tables S1 and S2). Hence these data characterize KDM2A/KDM2B as prominent PRC1-associated transcriptional repressors, in line with results of Gene Ontology analyses (Figure S3A).

**Figure 2:**
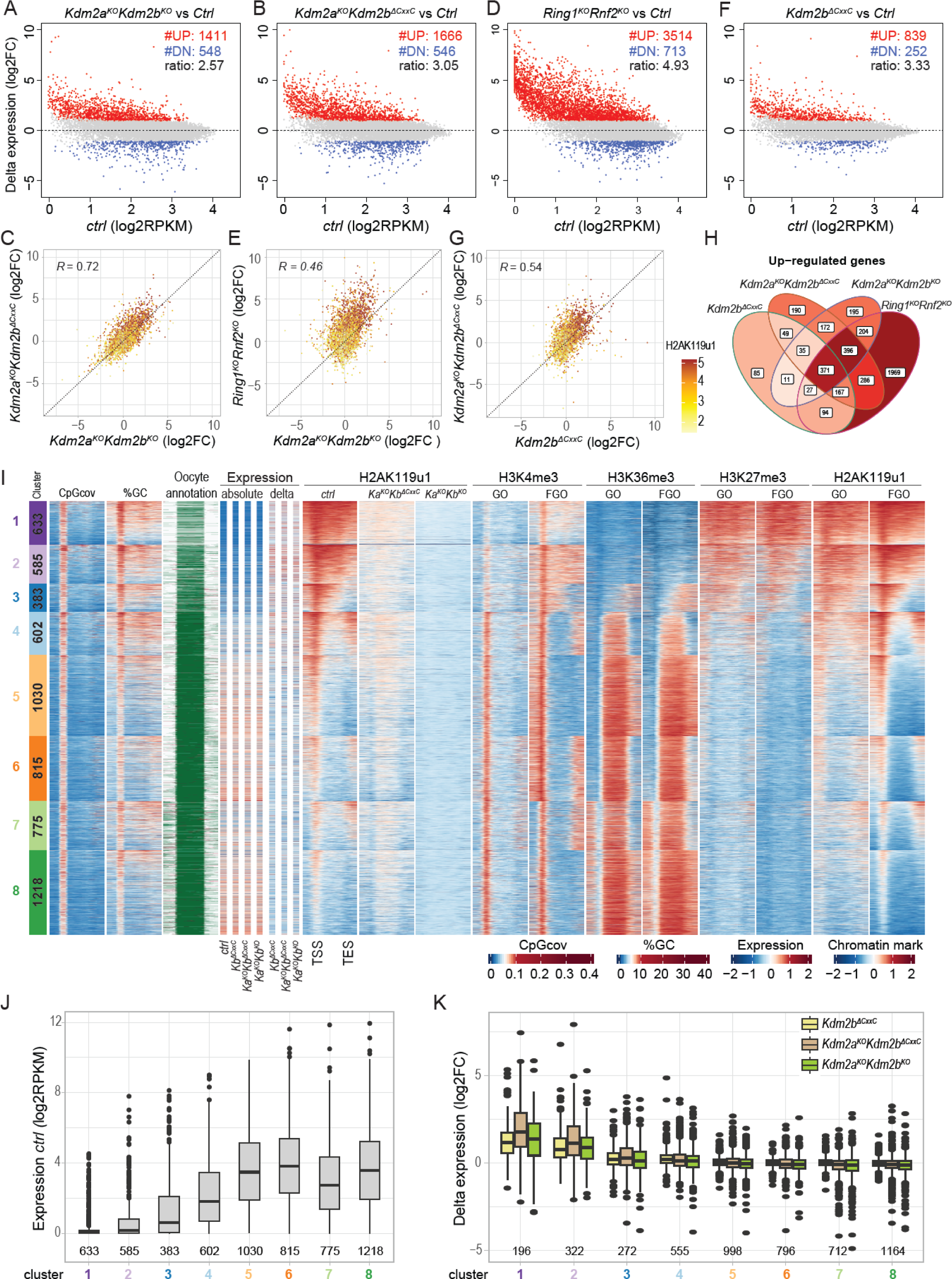
KDM2A/KDM2B regulate H2AK119u1 deposition and gene expression during oogenesis. **A.** MA-plot showing differential expression of *Kdm2a^KO^Kdm2b^KO^* over *ctrl* FGOs (log2 fold change (log2FC)) as a function of expression in *ctrl* FGOs (log2RPKM). #UP and #DN refer to numbers of genes more highly or lowly expressed in mutant versus *ctrl* FGOs (log2FC > 1.0; adj P-value < 0.05). Ratio refers to #UP genes over #DN genes. **B.** MA-plot as in Figure 2A, for *Kdm2a^KO^Kdm2b^ΔCxxC^* over *ctrl* FGOs. **C.** Scatter plot showing log2FC in expression of *Kdm2a^KO^Kdm2b^KO^* over *ctrl* FGOs versus *Kdm2a^KO^Kdm2b^ΔCxxC^* over *ctrl* FGOs. H2AK119u1 occupancy (log2) at promoters (-1500/+500 bps of TSS) is indicated by color scale ^36^. R indicates Pearson’s correlation coefficient. **D.** MA-plot as in Figure 2B, for *Ring1^KO^Rnf2^KO^* over *ctrl* FGOs. **E.** Scatter plot showing log2FC in expression of *Kdm2a^KO^Kdm2b^KO^* over *ctrl* FGOs versus *Ring1^KO^Rnf2^KO^* over *ctrl* FGOs. H2AK119u1 occupancy and R as in panel 2C. **F.** MA-plot as in Figure 2C, for *Kdm2b^ΔCxxC^* over *ctrl* FGOs. **G.** Scatter plot showing log2FC in expression of *Kdm2b^ΔCxxC^* over *ctrl* FGOs versus *Kdm2a^KO^Kdm2b^ΔCxxC^* over *ctrl* FGOs. H2AK119u1 occupancy and R as in panel 2C. **H.** Venn diagram showing numbers of genes up-regulated in *Kdm2b^ΔCxxC^*, *Kdm2a^KO^Kdm2b^ΔCxxC^*, *Kdm2a^KO^Kdm2b^KO^* and/or *Ring1^KO^Rnf2^KO^* FGOs. **I.** Heatmap displaying sequence composition, transcriptional and chromatin variables within CGI-promoter genes (5kb upstream, TSS, gene body, TES, and 5 kb downstream) grouped into 8 gene clusters by k-means clustering. Clusters 1 to 8 contain 633, 585, 383, 602, 1030, 815, 775 and 1218 genes. From left to right: CpG coverage; GC percentage; oocyte specific sense (green) and antisense (red) transcripts^40^; absolute RNA (scaled RPKM) in *ctrl*, *Kdm2b^ΔCxxC^*, *Kdm2a^KO^Kdm2b^ΔCxxC^* and *Kdm2a^KO^Kdm2b^KO^* FGOs; log2FC expression in *mutant* vs *ctrl* FGOs (delta); H2AK119u1 occupancy in FGOs of indicated genotypes; H3K4me3, H3K36me3, H3K27me3 and H2AK119u1 occupancies in wildtype GOs and FGOs^21,19,36^. All chromatin data are shown as Z-scores. Expression correlates with H3K4me3 promoter occupancy and H3K36me3 gene body occupancy while repression with broad H3K27me3 and H2AK119u1 occupancy, and low H3K4me3 promoter occupancy, particularly in GOs. **J.** Boxplot presenting RNA expression levels (in log2RPKM) per gene cluster in *ctrl* FGOs. Numbers of genes per cluster are indicated. **K.** Boxplot presenting log2FC in expression per gene cluster measured in various *mutant* relative to respective *ctrl* FGOs.

The comparable misexpression in *Kdm2a^KO^Kdm2b^KO^* and *Kdm2a^KO^Kdm2b^ΔCxxC^* FGOs suggests that binding of KDM2B to CGI-gene promoters via its CxxC domain is key for vPRC1-driven gene repression (Figure 2C). Moreover, of the 1666 genes upregulated in *Kdm2a^KO^Kdm2b^ΔCxxC^* FGOs, only 37% were upregulated in single *Kdm2b^ΔCxxC^* FGOs. Yet, another 41% were upregulated in *Ring^KO^Rnf2^KO^* FGO (Figures 2F, 2G, 2H, Tables S1 and S2). Together, contrasting the results in ESCs^29^, our data support the notion that KDM2A functions as a vPRC1 member in oocytes, like KDM2B.

### KDM2A/KDM2B recruit repressive PRC1 to chromatin

In FGOs, H2AK119u1 and H3K27me3 were previously reported to co-occupy broad genomic regions, while dual marking by H2AK119u1 and H3K4me3 were shown to label promoters of expressed genes^36,37,21,38^. To derive the syntax of DNA sequence and chromatin configurations underlying the gene regulatory function of KDM2A/KDM2B proteins, we selected CGI and non-CGI promoter genes (Figure S3C) and partitioned each gene group into eight clusters by k-means clustering, based on occupancy levels of H3K4me3^21^, H3K27me3^39^ and H2AK119u1^36^ at promoters and of H3K36me3^19^ along gene bodies measured in *wt* FGOs (Figures 2I and S3D). For CGI-promoter genes, this resulted in clusters 1-3 harboring Polycomb-regulated genes while clusters 4-8 contain genes that had been transcribed and hence accumulated H3K36me3 during oocyte growth (Figures 2I and 2J). We then integrated absolute and differential expression levels of *ctrl* and mutant FGOs into these clusters (Figures 2I-2K and S3D-S3F). We further measured local H2AK119u1 occupancies by CUT&RUN and observed an extensive to almost complete loss of the mark in all gene clusters in *Kdm2a^KO^Kdm2b^ΔCxxC^* and *Kdm2a^KO^Kdm2b^KO^* oocytes respectively, both relative to *ctrl* FGOs (Figures 2I and S3D). CGI-promoter genes in clusters 1, 2 and some in 3, characterized by extensive H2AK119u1 and H3K27me3, lack of H3K36me3 and low to no expression in *wt* GOs and FGOs were primarily upregulated in *Kdm2b^ΔCxxC^*, *Kdm2a^KO^Kdm2b^ΔCxxC^* and *Kdm2a^KO^Kdm2b^KO^* oocytes. In contrast, expressed genes in clusters 4-8 with moderate to low H2AK119u1 but high H3K4me3 levels at their promoters barely responded to KDM2A/KDM2B mutations (Figures 2I-2K). We observed a similar chromatin logic for non-CGI promoter genes, for which H2AK119u1 and H3K27me3 co-occupancy in *wt* oocytes relates to CpG/GC density (Figures S3D-S3F). Hence, while the KDM2A/KDM2B proteins control H2AK119u1 deposition at gene promoters throughout the genome, they only repress genes extensively labeled with H2AK119u1, and also marked by PRC2-mediated H3K27me3.

### KDM2A/KDM2B restrict H3K36me2 and DNAme deposition in genes

Besides their role in vPRC1 recruitment to DNA, KDM2A/KDM2B demethylate mono- and di-methylation of H3K36 (H3K36me1/2)^28^. To test their catalytic function during oogenesis, we performed IF staining of GOs and measured an increase in H3K36me2 in *Kdm2a^KO^Kdm2b^ΔCxxC^* mutants. The increase was more significant in *Kdm2a^KO^Kdm2b^KO^* oocytes, which completely lack KDM2A/KDM2B demethylase activity (Figures 1A and 3A). In contrast, we observed only minor increases in H3K36me3 in mutant GOs, which may relate to the slightly enhanced gene expression that we measured in such mutant FGOs (Figures 2K, S3F and 3B)^19^.

**Figure 3:**
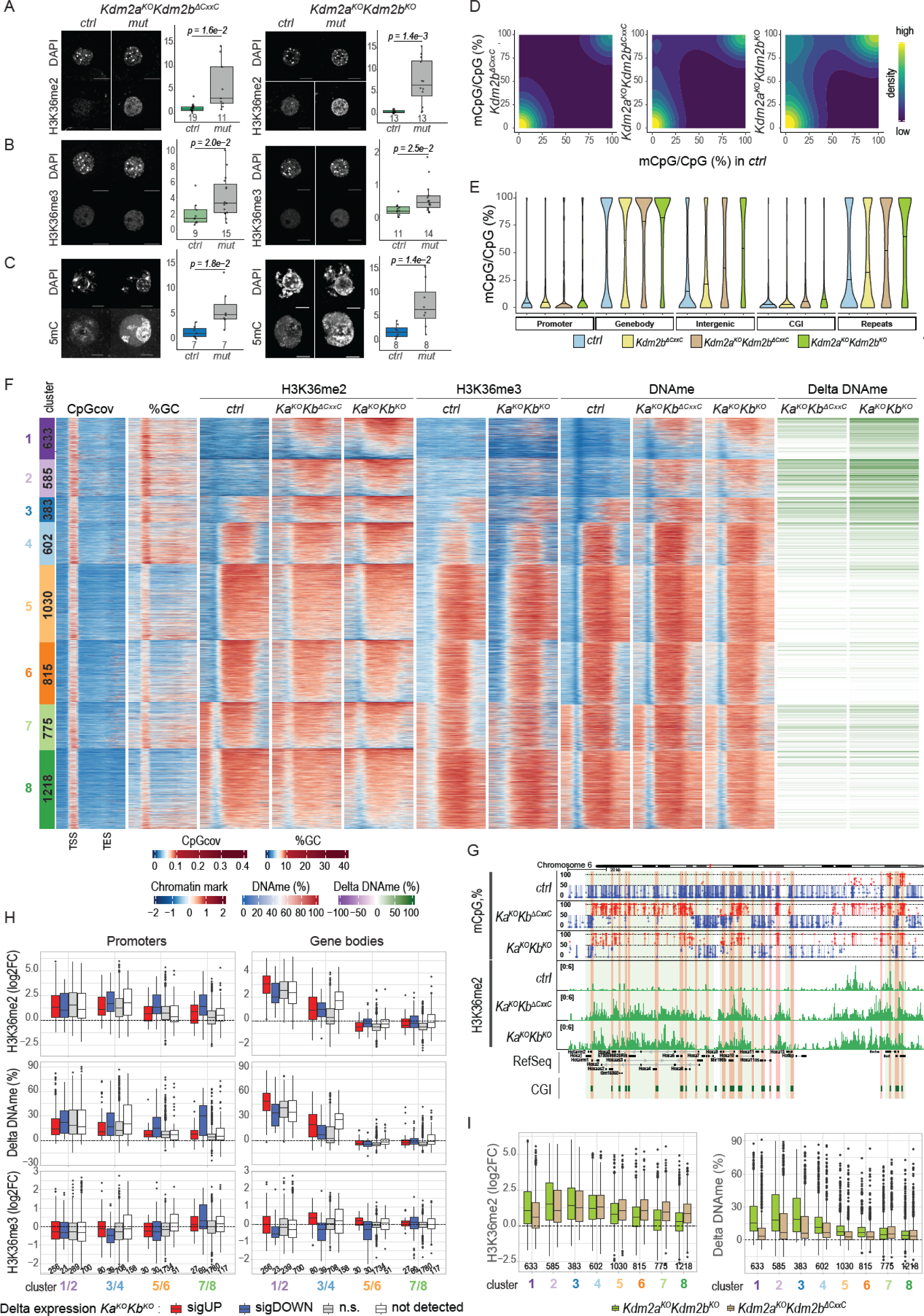
KDM2A/KDM2B restrict H3K36me2 and DNAme accumulation at promoters and along genes in oocytes. **A. B.** Immunofluorescence staining and quantification of H3K36me2 (**A**) and H3K36me3 (**B**) in GOs of *Kdm2a^KO^Kdm2b^ΔCxxC^*, *Kdm2a^KO^Kdm2b^KO^* and *ctrl* genotypes. Number of analyzed oocytes are indicated. P-values according to two-sided student’s t-test. **C.** Immunofluorescence staining and quantification of 5mC in FGOs of *Kdm2a^KO^Kdm2b^ΔCxxC^, Kdm2a^KO^Kdm2b^KO^* and *ctrl* genotypes. Number of analyzed oocytes are indicated. P-values according to two-sided student’s t-test. **D.** 2D-density plots displaying distributions of CpGs according to their mean methylation levels (mCpG/CpG in %) in *Kdm2b^ΔCxxC^, Kdm2a^KO^Kdm2b^ΔCxxC^* and *Kdm2a^KO^Kdm2b^KO^* FGOs versus *ctrl* FGOs. **E.** Violin plot showing distribution of mCpG/CpG (%) values at different genome elements in FGOs of indicated genotypes (*ctrl, Kdm2b^ΔCxxC^*, *Kdm2a^KO^Kdm2b^ΔCxxC^* and *Kdm2a^KO^Kdm2b^KO^*). **F.** Heatmap displaying sequence composition and chromatin variables within 8 CGI-promoter gene clusters in FGOs, identical to those described in Figure 2I. From left to right: CpG coverage; GC percentage; H3K36me2, H3K36me3 and DNAme in FGOs of indicated genotypes; differential (Delta) DNAme at CGI promoters in *Kdm2a^KO^Kdm2b^KO^* and *Kdm2a^KO^Kdm2b^ΔCxxC^* FGOs. **G.** Genomic snapshot of the *Hoxa* and *Evx1* gene cluster gaining DNAme and H3K36me2 at CGI promoters (highlighted in orange) and gene bodies in *Kdm2a^KO^Kdm2b^ΔCxxC^* and *Kdm2a^KO^Kdm2b^KO^* FGOs. DNA methylation and H3K36me2 are indicated. **H.** Boxplots displaying differential H3K36me2, DNAme and H3K36me3 at CGI-gene promoters (- 1500/+500 bps of TSS) or gene bodies (+500 bps of TSS to TES) of genes belonging to merged gene clusters for which expression is either upregulated, downregulated, not changed or not detected in *Kdm2a^KO^Kdm2b^KO^* FGOs relative to *ctrl* FGOs. Numbers of genes per condition are indicated. **I.** Boxplots displaying differential H3K36me2 and DNAme at promoters (-1500/+500 bps of TSS) of all genes belonging to the 8 CGI-promoter gene clusters in *Kdm2a^KO^Kdm2b^KO^* and *Kdm2a^KO^Kdm2b^ΔCxxC^* FGOs relative to *ctrl* FGOs, as indicated. Numbers of genes per cluster are indicated.

Given the role of *Kdm2b* in preventing DNAme at Polycomb-regulated CGIs in ESCs^25^, we next assessed DNAme by IF, revealing a vast increase in 5mC levels in both mutants (Figure 3C). To determine which genomic regions gain DNAme, we performed whole genome bisulfite sequencing (WGBS) (Figures S4A-S4C). We measured considerable gains in methylated CpG dinucleotides from a global average of 35.8% mCpGs/CpGs in controls to 40.7% in *Kdm2b^ΔCxxC^*, 51.5% in *Kdm2a^KO^Kdm2b^ΔCxxC^* and 58.8% in *Kdm2a^KO^Kdm2b^KO^* FGOs (Figures 3D and S4C). DNAme gains were not confined to Polycomb-controlled promoter regions only, as reported previously for *Kdm2b* deficient ESCs^25^ but also extended widely into gene bodies, intergenic regions, and endogenous repetitive elements (ERVs) (Figures 3E and 3F) as exemplified for the *Hoxa* gene cluster (Figure 3G). Importantly non-Polycomb controlled promoter CGIs acquired aberrant DNAme as well (Figures 3F and 3H).

To understand the mechanism(s) underlying the (differential) increases in DNAme between mutants, we aimed to compare the genome-wide distribution of DNAme to those of H3K36me2 and H3K36me3. We therefore performed CUT&RUN analyses for H3K36me2 in *ctrl*, *Kdm2a^KO^Kdm2b^ΔCxxC^* and *Kdm2a^KO^Kdm2b^KO^* FGOs as well as for H3K36me3 in *ctrl* and *Kdm2a^KO^Kdm2b^KO^* FGOs (Figure S4D). We first analyzed changes at intragenic and promoter sequences (Figures 3F-3I) and then at intergenic regions (Figure 4). In *ctrl* oocytes, we observed a strong co-occupancy of H3K36me2, H3K36me3 and DNAme within gene bodies of transcribed genes from CGI promoters (clusters 4-8), as reported previously for H3K36me3 and DNAme (Figure 3F displaying identical gene clusters shown in Figure 2l)^20,19^. Regions upstream (clusters 7, 8) and downstream (clusters 5, 8) of expressed genes showed similar marking, likely reflecting initiation of gene transcription from upstream located ERVs^40^ and transcriptional run-through beyond the annotated transcriptional end site (TES), respectively (Figure 3F).

**Figure 4:**
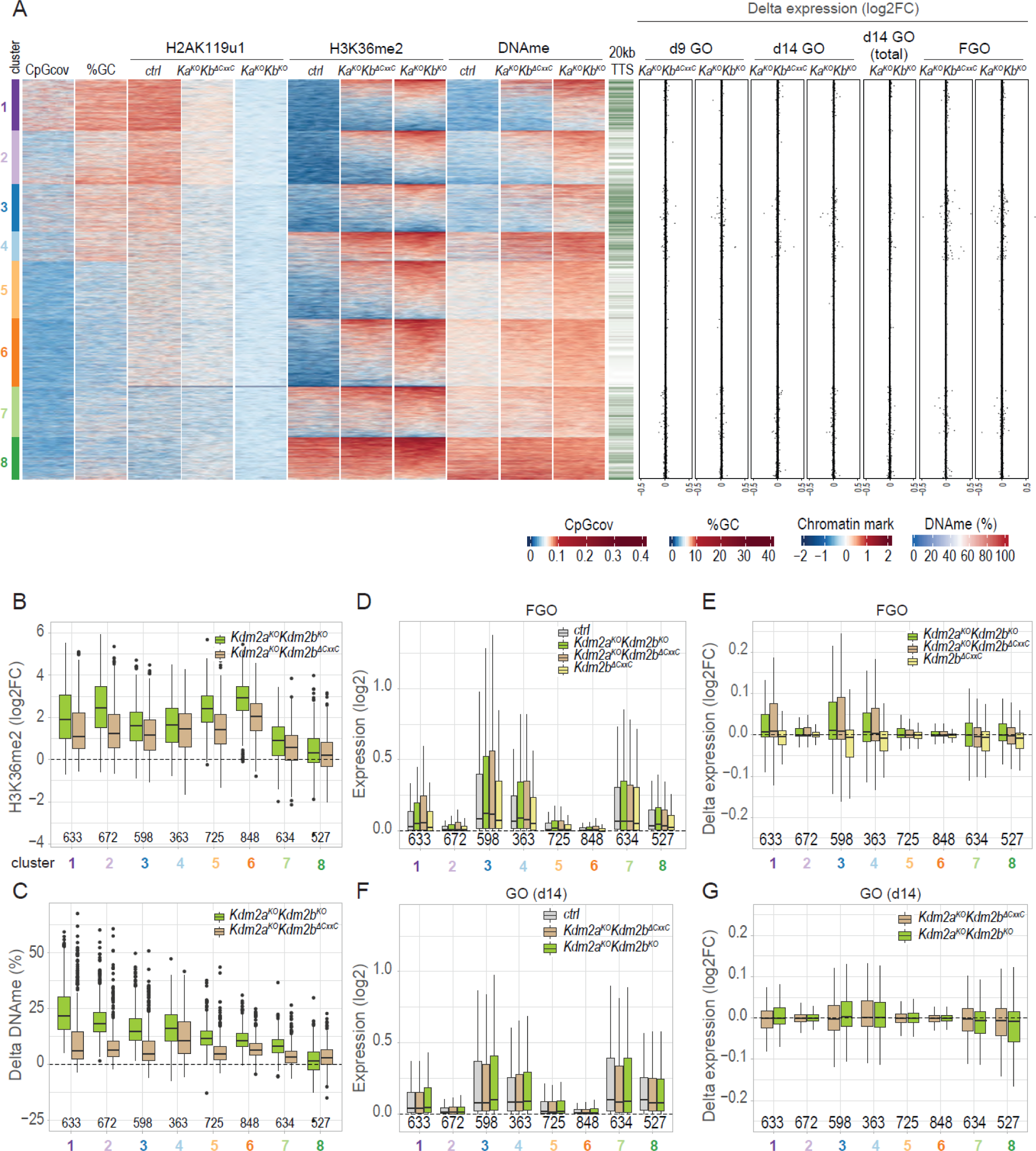
H3K36me2 and DNAme accumulate throughout the genome of *Kdm2a*/*Kdm2b* deficient oocytes, independently of transcription. **A.** Heatmap displaying chromatin and transcriptional variables within 8 clusters of 10 kb intergenic regions in oocytes. Data is displayed in 20 neighboring 500 bp bins. From left to right: CpG coverage; GC percentage; H2AK119u1, H3K36me2 and DNAme in *ctrl, Kdm2a^KO^Kdm2b^KO^* and *Kdm2a^KO^Kdm2b^ΔCxxC^* FGOs; presence of annotated TTS in 20 kb flanking regions, that could be compatible with run-through transcription through the window; log2FC in expression between indicated *mutant* and *ctrl* genotypes in day9 GOs; in day14 GOs; in random primed (total) day14 GOs and in FGOs. All RNA expression data is based on polyA-primed RNA capture and Smart-seq2 library generation, if not indicated otherwise. **B. C.** Boxplots displaying differential H3K36me2 (**B**) and DNAme (**C**) for 10 kb intergenic regions in 8 clusters in *Kdm2a^KO^Kdm2b^KO^* or *Kdm2a^KO^Kdm2b^ΔCxxC^* FGOs relative to *ctrl* FGOs, as indicated. Numbers of regions per cluster are indicated. **D.** Boxplot displaying absolute expression for 10 kb intergenic regions in 8 clusters in *ctrl*, *Kdm2b^ΔCxxC^*, *Kdm2a^KO^Kdm2b^ΔCxxC^* and *Kdm2a^KO^Kdm2b^KO^* FGOs, as indicated. Numbers of regions per cluster are indicated. **E.** Boxplot displaying log2FC expression for 10 kb intergenic regions in 8 clusters in *Kdm2b^ΔCxxC^*, *Kdm2a^KO^Kdm2b^ΔCxxC^* and *Kdm2a^KO^Kdm2b^KO^* FGOs relative to *ctrl* FGOs, as indicated. Numbers of regions per cluster are indicated. **F.** Boxplot displaying absolute expression for 10 kb intergenic regions in 8 clusters in *ctrl* and *Kdm2a^KO^Kdm2b^ΔCxxC^* and *Kdm2a^KO^Kdm2b^KO^* GOs at day14. Numbers of regions per cluster are indicated. **G.** Boxplot displaying log2FC expression for 10 kb intergenic regions in 8 clusters in *Kdm2a^KO^Kdm2b^ΔCxxC^* and *Kdm2a^KO^Kdm2b^KO^* GOs relative to *ctrl* GO GOs at day14. Numbers of regions per cluster are indicated.

In *Kdm2a^KO^Kdm2b^ΔCxxC^* and *Kdm2a^KO^Kdm2b^KO^* FGOs, we measured major gains in H3K36me2 and DNAme along gene bodies of cluster 1-3 genes compared to *ctrl* oocytes, with most genes reaching high levels characteristic of robustly expressed genes as those in clusters 5-8 (Figure 3F). In contrast, H3K36me3 levels remained largely unchanged along gene bodies of cluster 1-3 genes (Figure 3F) suggesting that the rather moderate increases in expression upon *Kdm2a*/*Kdm2b* deficiency (Figures 2I and 2K) are insufficient for robust *Setd2*-dependent H3K36me3 deposition^19^. Hence, it is unlikely that aberrant DNAme acquisition at cluster 1-3 genes in mutant FGOs was instructed by H3K36me3 (Figure 3F). Quantitative enrichment analysis revealed indeed strong gains of both H3K36me2 and DNAme, but not of H3K36me3 in gene bodies of CGI- and non-CGI-promoter genes (Figures 3H, S4E and data not shown). Importantly, H3K36me2 and DNAme were also increased at genes with unchanged or down-regulated expression, or with non-detectable expression in *Kdm2a*/*Kdm2b* FGOs (Figure 3H). Further, gains in H3K36me2 and DNAme were more pronounced along cluster 1-3 genes of *Kdm2a^KO^Kdm2b^KO^* compared to *Kdm2a^KO^Kdm2b^ΔCxxC^* oocytes (Figures 3H, 3I and S4E) and even occurred at regions upstream and/or downstream of genes (Figure 3F). Together, these data argue that the gain in H3K36me2 along gene bodies is not linked to transcription but is resulting from the loss of H3K36me2 demethylase activity by KDM2A/KDM2B. The DNAme data is further in line with that DNMT3A is recruited to chromatin via its PWWP domain binding to elevated H3K36me2 levels, catalyzing *de novo* DNAme.

### KDM2A/KDM2B control H3K36me2 and DNAme at CGI promoters, irrespective of PRC1 regulation

In addition to gene bodies, we measured increased H3K36me2 and DNAme at promoter regions of CGI- and non-CGI-promoters genes marked by H2AK119u1/H3K27me3 in *wt* oocytes (Figures 2I, 3F, 3H, S3D and S4E). Remarkedly, even at CGI-promoter genes belonging to clusters 4-8, which are not or only weakly marked by H2AK119u1, lack H3K27me3 at their promoters, and are expressed in *ctrl* oocytes, we measured significant gains in H3K36me2 and DNAme in both types of mutant oocytes, particularly for those genes displaying decreased expression (Figures 2I, 3F, 3H and 3I). These results differ from the reported gain in DNAme in *Kdm2b*-deficient ESCs, occurring exclusively at CGI promoters controlled by PRC1^25^. Instead, consistent with KDM2A and KDM2B localizing to all CGI-promoters in ESCs irrespective of their chromatin status^29,31^, our data point to a wide-spread catalytic H3K36me2 demethylating function of KDM2A and KDM2B *in vivo*, protecting PRC1- and non-PRC1-controlled CGI promoters from gaining H3K36me2 and DNAme, and from becoming aberrantly repressed.

### KDM2A/KDM2B prevent deposition of H3K36me2 and DNAme genome wide

To investigate whether non-genic regions respond similarly, we partitioned 10kb-sized intergenic sequences in 8 clusters based on H3K4me3 ^21^, H3K36me3 ^19^, H3K27me3 ^39^ and H2AK119u1^36^ occupancy in *wt* FGOs (Figure S5A) and then integrated changes in H3K36me2, H3K36me3 and DNAme levels in mutant versus *ctrl* FGOs as well (Figure 4A). As for genes, GC-dense regions in clusters 1-3, broadly marked by H2AK119u1 and H3K27me3, were devoid of H3K36me2, H3K36me3 and DNAme in *ctrl* FGOs (Figures 4A and S5A). These regions gained H3K36me2 and DNAme moderately to majorly in *Kdm2a^KO^Kdm2b^ΔCxxC^* and *Kdm2a^KO^Kdm2b^KO^* FGOs, respectively (Figures 4A-4C). Even GC-poor regions in clusters 5-7, marked by low H2AK119u1 levels in *ctrl* GOs gained H3K36me2 and DNAme, particularly again in *Kdm2a^KO^Kdm2b^KO^* FGOs (Figures 4A-4C). Hence, these data point to a prominent genome-wide role, beyond CGIs, for KDM2A/KDM2B proteins in maintaining H3K36me2 levels low during oocyte development.

We did not measure consistent upregulation of expression within the 10kb-nor within flanking regions having gained aberrant H3K36me2 in mutant FGOs (Figures 4A, 4D and 4E). To investigate possible changes in expression during oocyte growth, we profiled GOs at day 9 and 14 of development using poly-A and random primed RNA-seq. approaches. Again, we did not measure consistent transcript level upregulation in regions gaining aberrant H3K36me2 (Figures 4F, 4G and S5D-S5G). In summary, based on these genic and intergenic assessments, our data support a wide-spread transcription-independent deposition of H3K36me2 during oocyte growth that is efficiently counteracted by KDM2A/KDM2B proteins.

### H3K4me3 in growing oocytes prevents atypical DNAme acquisition

To identify chromatin features underlying the aberrant *Dnmt3a*-mediated DNA methylation at CpG islands, we performed regularized linear regression analysis for predicting DNAme, assuming additive effects of different sequence and chromatin input parameters in GOs and FGOs. First, almost 83% of the variation in basal CGI DNAme in *ctrl* FGOs could be explained, demonstrating the suitability of the approach (Figures S6A-S6C). In keeping with previous reports^11,20, 19^, H3K36me2 and H3K36me3 contributed positively, while H3K4me3 in *wt* FGOs represented the key negative predictor to CGI DNAme (Figures 5A, S6B and S6C)^21^.

**Figure 5:**
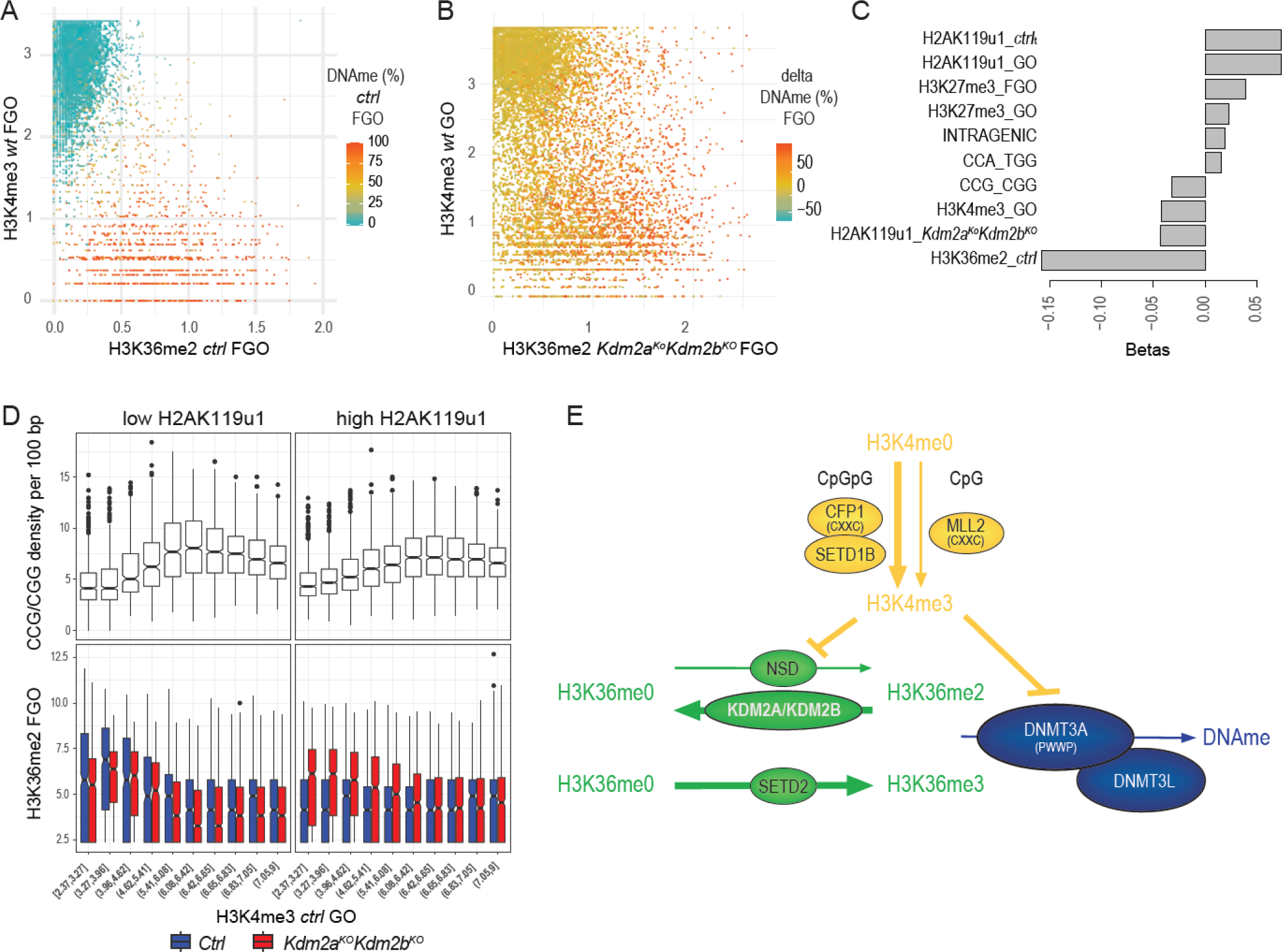
Low H3K4me3 in GOs is permissive for H3K36me2 and DNAme acquisition during oocyte growth. **A.** Scatter plot showing correlations between H3K36me2 and H3K4me3 occupancies, and DNAme (%) at promoter and intragenic CGIs in *ctrl* and *wt* FGOs. **B.** Scatter plot showing the difference in DNAme (%) at promoter and intragenic CGIs between *Kdm2a^KO^Kdm2b^KO^* over *ctrl* FGOs in relation to H3K36me2 occupancy in *Kdm2a^KO^Kdm2b^KO^* FGOs and H3K4me3 occupancy in *wt* GOs. **C.** Barplot showing linear regression coefficients of chromatin features and triplet nucleotide sequences contributing to predicting differential H3K36me2 occupancy at CGIs in *Kdm2a^KO^Kdm2b^KO^* over *ctrl* FGOs (*R^2^*=0.343). **D.** Boxplots representing frequencies of CCG/CGG trinucleotides per 100bp (top panels) and enrichments of H3K36me2 in *ctrl* (blue) and *Kdm2a^KO^Kdm2b^KO^* (red) FGOs (bottom panels) at 533bp regions surrounding CGI centers for different bins of H3K4me3 occupancies in GOs for CGIs with low (left panels) and high (right panels) H2AK119u1 levels. **E.** Cartoon illustrating DNA-sequence dependency as well as hierarchies and dependencies between different histone modifying enzymes and DNA methyltransferases in regulating H3K36 di-/tri-methylation and *de novo* DNA methylation.

When employing chromatin features of *ctrl* oocytes to predict aberrant gains of DNAme in *Kdm2a^KO^Kdm2b^KO^* versus *ctrl* FGOs, only 33% of variation could be explained (data not shown). When including H3K36me2, H3K36me3 and H2AK119u1 occupancies in *Kdm2a^KO^Kdm2b^KO^* FGOs into the regression analysis as well, 64% of the change in DNAme at CGIs could be accounted for, with H3K36me2 contributing positively and residual H2AK119u1 in *Kdm2a^KO^Kdm2b^KO^* FGOs contributing negatively (Figures S6D-S6F). Remarkably, H3K4me3 occupancy as measured in *wt* GOs, but not FGOs, was a negative correlate (Figure S6F). This suggests that CGIs are permissive for aberrant *de novo* DNAme only when H3K4me3 occupancy levels are low early during oocyte growth (Figure 5B)^20^. In summary, our data is consistent with high H3K36me2 and low H3K4me3 occupancy serving instructive and permissive functions, respectively, for acquisition of atypical DNAme at CGIs in *Kdm2a^KO^Kdm2b^KO^* and *Kdm2a^KO^Kdm2b^ΔCxxC^* growing oocytes.

### Sequence and chromatin features define aberrant H3K36me2 acquisition at CGIs

Our analysis clearly shows that atypical DNAme is established downstream of aberrant H3K36me2. In turn, the globally increased H3K36me2 occupancy in *Kdm2a^KO^Kdm2b^KO^* FGOs likely reflects widespread H3K36 methyltransferase activity in GOs. Nonetheless, clustering analysis of CGIs indicated that H3K36me2 occupancy in *Kdm2a^KO^Kdm2b^KO^* FGOs was only increased at ∼30% of such elements that are normally controlled by PRC1 (data not shown). To start addressing the regulatory complexity underlying specification of atypical H3K36me2 occupancy at CGIs, we performed regularized linear regression analysis for H3K36me2 itself (Figures 5C and S6G-S6H). This showed that aberrant H3K36me2 in *Kdm2a^KO^Kdm2b^KO^* FGOs is indeed deposited at CGIs normally marked by H2AK119u1 and to some extent by H3K27me3 in *wt* FGOs. In line, residual H2AK119u1 in *Kdm2a^KO^Kdm2b^KO^* FGOs associated negatively with atypical H3K36me2, a finding supported by biochemical studies demonstrating robust inhibition of all NSD and SETD2 KMTs by nucleosomal H2AK119u1^41,42^. Moreover, as for DNA methylation, H3K4me3 in GOs is a negative predictor for atypical H3K36me2 occupancy, suggesting that H3K36 KMT function *in vivo* is inhibited by H3K4me3. This latter finding is in line with biochemical results for NSD3^42^.

We next aimed at better understanding the rationale underlying the heterogeneity in H3K4me3 occupancy among CGIs in GOs and its possible negative impact on H3K36me2 and DNAme acquisition in *Kdm2a^KO^Kdm2b^KO^* GOs. At most CGI promoters in mouse oocytes, H3K4me3 is robustly catalyzed by the SETD1B KMT in conjunction with the CFP1 (CXXC1) cofactor binding preferentially to CpGpG trinucleotides and reading out H3K4me3 as well^43,39,44,45^. At other promoters, intra- and intergenic sites, H3K4me3 is deposited by MLL2, which is recruited via its CXXC domain to CpG dinucleotides having no preference for adjacent bases^21,46^. When including trinucleotide frequencies into our modeling, we identified a robust negative contribution of CpGpG trinucleotides to atypical H3K36me2 in *Kdm2a^KO^Kdm2b^KO^* FGOs (Figure 5C), suggesting that SETD1B/CFP1 catalyzing robust H3K4me3 may be a major barrier to NSD and/or SETD2-mediated catalysis at promoter CGIs in oocytes. Consistently, CGIs with low H3K4me3 in GOs are characterized by low CpGpG densities and harbor higher H3K36me2 in FGOs when H2AK119u1 levels are reduced (Figures 5D, S6I and S6J). In summary, our analyses reveal that a selective subset of CGIs with low frequency of CpGpG trinucleotides is particularly vulnerable towards aberrant accumulation of H3K36me2 and DNAme (Figure 5E)^42^.

### Aberrant maternal DNAme impairs pre-implantation development

To investigate the impact of the ∼50% increase in DNAme levels (Figure 3D) in the maternal genome of *Kdm2aKdm2b* mutant oocytes for embryonic development, we first investigated its stability upon fertilization by performing IF analysis on zygotes. Impressively, contrasting to the global loss of DNAme of the sperm genome, aberrant methylation in the maternal genome originating from *Kdm2a^KO^Kdm2b^KO^* oocytes did not become distinctively reprogrammed upon fertilization (Figure 6A), in keeping with parent-of-origin specific epigenetic reprogramming of regular DNAme^3^. In line with less extensive gains in aberrant DNAme in *Kdm2a^KO^Kdm2b^ΔCxxC^* (than *Kdm2a^KO^Kdm2b^KO^*) FGOs, we did not observe a significant difference in global maternal DNAme levels in *Kdm2a^matKO^Kdm2b^matΔCxxC^* zygotes versus *ctrl* zygotes (Figure 6B). We next addressed whether the aberrant maternal methylome drives the low developmental competence of both types of *Kdm2aKdm2b* mutant oocytes. To do so, we prevented the establishment of DNAme in *Kdm2a^KO^Kdm2b^KO^* and *Kdm2a^KO^Kdm2b^ΔCxxC^* GOs by conditionally mutating *Dnmt3a* function^12^. Critically, embryos maternally triple deficient developed into blastocyst stage embryos equally efficiently as control embryos. Thus, maternal *Dnmt3a* deficiency elicited complete suppression of the progressive embryonic lethality seen for both maternal *Kdm2aKdm2b* mutants (compare Figure 1B to Figures 6C and 6D). In contrast, deficiency of the DNMT1 enzyme, normally contributing together with UHRF1 to *de novo* DNAme at certain inactive regions in late GOs and maintaining DNAme during pre-implantation development^47,48,49^, did not rescue the embryonic lethality caused by *Kdm2aKdm2b* deficiency in oocytes (Figures 1B, 6C and 6D). Hence, these data indicate that beyond regulating vPRC1-mediated gene repression, the prime intergenerational function of KDM2A and KDM2B in oocytes is confining the targeting of DNMT3A and *de novo* DNAme catalysis, by preventing H3K36me2 accumulation throughout the genome.

**Figure 6:**
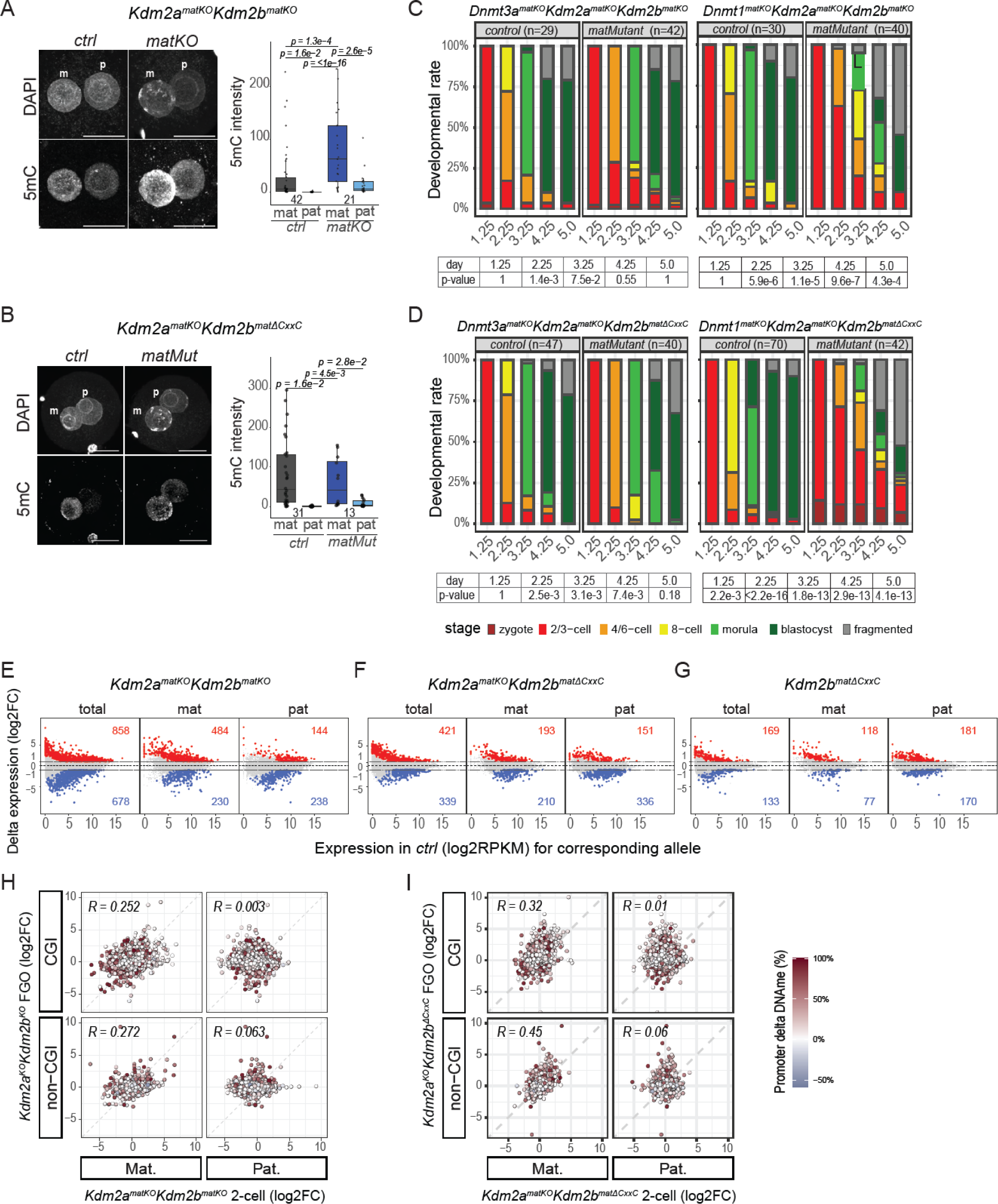
Aberrant maternal DNA methylation impairs pre-implantation development. **A. B.** Immunofluorescence staining and quantification of 5mC in maternal (mat) and paternal (pat) pronuclei of *Kdm2a^matKO^Kdm2b^matKO^* (**A**), *Kdm2a^matKO^Kdm2b^matΔCxxC^* (**B**) and respective *ctrl* late zygotes. Numbers of analyzed embryos oocytes are indicated. P-values according to Tukey’s HSD test. **C. D.** Developmental progression rates of pre-implantation embryos at embryonic day e1.25, e2.25, e3.25, e4.25 and e5.0, generated by fertilizing *Dnmt3a^KO^Kdm2a^KO^Kdm2b^KO^*, *Dnmt1^KO^Kdm2a^KO^Kdm2b^KO^* (**C**), *Dnmt3a^KO^Kdm2a^KO^Kdm2b^ΔCxxC^*, *Dnmt1^KO^Kdm2a^KO^Kdm2b^ΔCxxC^* (**D**) triple knock-out and respective *ctrl* oocytes with *wt* sperm and cultured over 5 days *in vitro*. Numbers of analyzed embryos are indicated. P-values according to Fisher’s exact test. See also Figure 1B. **E. F. G.** MA-plots showing differential expression in *Kdm2a^matKO^Kdm2b^matKO^* (**E**), *Kdm2a^matKO^Kdm2b^matΔCxxC^* (**F**) and *Kdm2b^matΔCxxC^* (**G**) over respective *ctrl* 2-cell embryos (log2FC) as a function of expression in *ctrl* 2-cell embryos (log2RPKM) for all sequencing reads (total) and for those mapping to the mat or pat genomes based on strain specific SNPs. Genes, Up or Down regulated in mutants, are indicated in red and blue (|log2FC| > 1.0; adj P-value < 0.05). **H.** Scatter plots showing log2FC expression of *Kdm2a^matKO^Kdm2b^matKO^* over *ctrl* 2-cell embryos versus log2FC expression of *Kdm2a^KO^Kdm2b^KO^* over *ctrl* FGOs for mat and pat specific expression of CGI- and non-CGI promoter genes. Differential promoter methylation in *Kdm2a^KO^Kdm2b^KO^* over *ctrl* FGOs is indicated by color (in %). **I.** Scatter plots as in Figure 6H, for *Kdm2a^matKO^Kdm2b^matΔCxxC^* 2-cell embryos and *Kdm2a^KO^Kdm2b^ΔCxxC^* FGOs relative to respective *ctrl* samples.

### Aberrant DNAme impairs gene expression in oocytes and early embryos

To assess the impact of maternal *Kdm2a*/*Kdm2b* deficiency on gene regulation in embryos, we profiled differential expression in *Kdm2a^matKO^Kdm2b^matKO^* versus *ctrl* two-cell embryos sired by JF1/Ms males allowing assessment of parental allelic specific expression (Figures S1B and S7A-S7D). Hundreds of genes were mis-regulated, with clear allele specific responses (Figure 6E, Tables S4 and S5). Comparative expression analyses between oocytes and 2-cell embryos revealed positive correlations for maternal differentially expressed alleles but not for those of paternal origin (Figure 6H). 36% of 157 maternally expressed CGI-promoter genes that were down-regulated in mutant 2-cell embryos were hypermethylated (>50%) at their promoters in *Kdm2a^KO^Kdm2b^KO^* oocytes (Figure 6H). Surprisingly, multiple methylated CGIs displayed up-regulated gene expression, especially in mutant FGOs (Figure 6H). We observed comparable transcriptional and methylation responses in *Kdm2a^matKO^Kdm2b^matΔCxxC^* and to a lesser degree in single mutant *Kdm2b^matΔCxxC^* samples (Figures 6F, 6G and 6I).

To investigate in more detail the mechanistic relationship between aberrant DNAme and transcription in the different (maternally) mutant oocytes and two-cell embryos, we first grouped CGIs located within UCSC-defined promoters in 7 clusters according to their DNAme status in *ctrl* FGOs, 3 types of mutant FGOs and *wt* sperm (Figure 7A). In clusters 1 to 4, almost 1500 promoter CGIs had gained extensive aberrant DNAme in *Kdm2aKdm2b* mutant FGOs. Further, over 1250 promoter CGIs of cluster 5 genes gained aberrant DNA to moderate levels (<50%), particularly in *Kdm2a^KO^Kdm2b^KO^* oocytes. Consistent with their identity as PRC1-target genes, GO-term analysis shows that many genes of cluster 1-5 CGI-promoters serve important functions during development (Figure 7B and Table S6). In contrast, cluster 6 contained CGIs that are highly methylated in *ctrl* and mutant FGOs. These CGIs likely localize within gene bodies transcribed from alternative upstream oocyte-specific promoters, despite their UCSC-classification as promoters based on somatic transcriptional data. In accord, these CGIs are marked by low H3K4me3 and high H3K36me3 occupancy levels, and are characterized by high transcript levels upstream of the CGI in *wt* FGOs (Figures S7E-S7G). Cluster 7 CGIs were either unmethylated or harbored only low DNAme levels in any genotype. Notably, none of the UCSC-annotated CGIs were substantially methylated in mature spermatozoa, indicating important differences between oocyte and sperm methylomes^50^ (Figure 7A).

**Figure 7:**
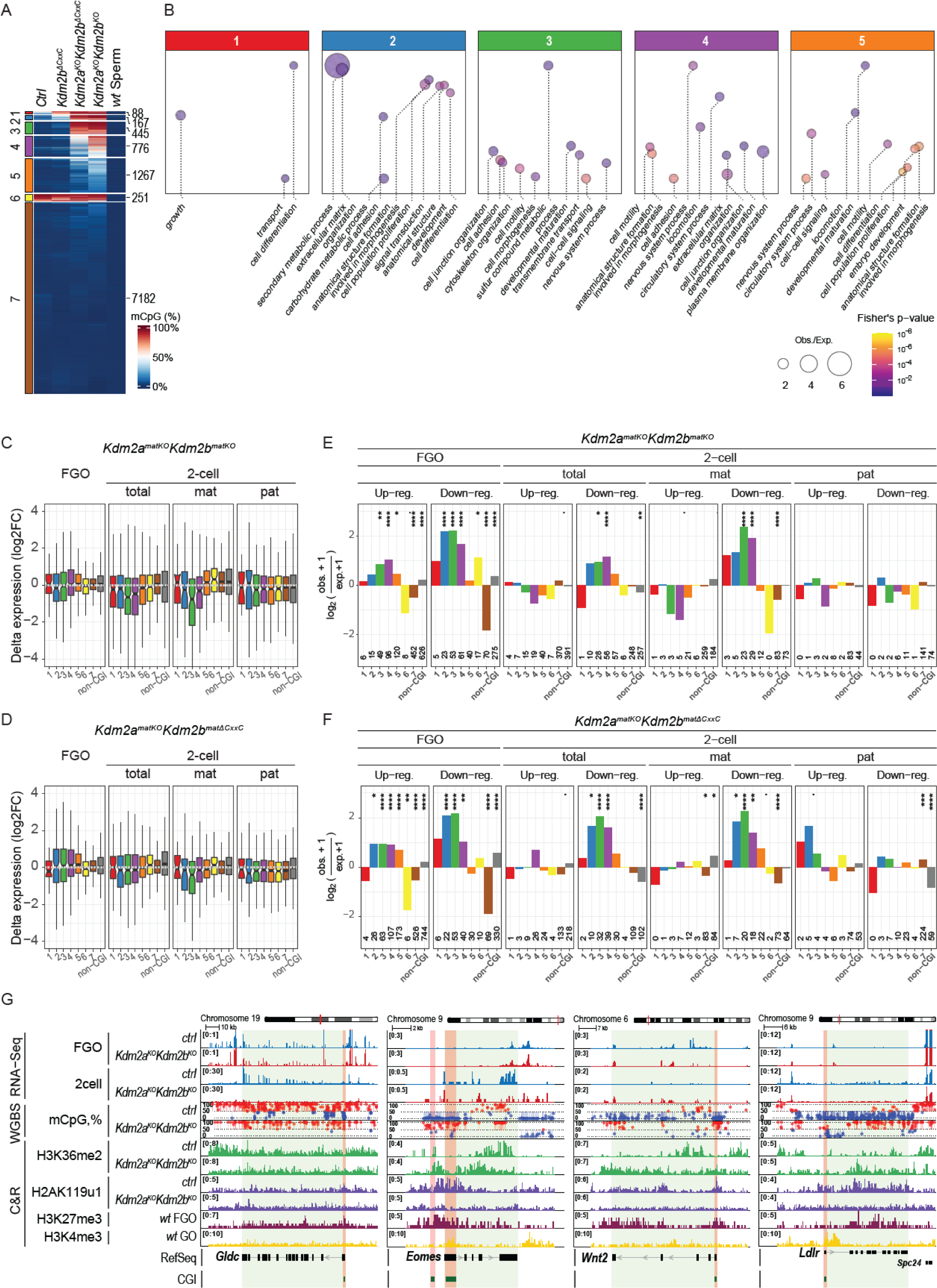
Genes marked by aberrant maternal promoter DNAme are repressed in early embryos. **A.** Heatmap showing clustering of CGI-promoter genes (based on UCSC annotation) according to absolute promoter methylation levels in *ctrl*, *Kdm2b^ΔCxxC^*, *Kdm2a^KO^Kdm2b^ΔCxxC^* and *Kdm2a^KO^Kdm2b^KO^* FGOs and in *wt* sperm^50^. Numbers of clustered genes are indicated. **B.** Gene Ontology enrichment analysis for genes associated to CGIs belonging to DNAme clusters 1-5 as defined in Figure 7A. Bubbles representing GO terms are scaled according to enrichments, colored according to statistical significance and positioned relative to one another to reflect similarities between significantly affected genes with corresponding GO terms. **C.** Boxplot showing log2FC expression of different clusters of CGI-promoter and all non-CGI-promoter genes in *Kdm2a^KO^Kdm2b^KO^* over *ctrl* FGOs and in *Kdm2a^matKO^Kdm2b^matKO^* over *ctrl* 2-cell embryos according to all, mat and pat specific sequencing reads. **D.** Boxplot as in Figure 7C, for *Kdm2a^KO^Kdm2b^ΔCxxC^* FGOs and *Kdm2a^matKO^Kdm2b^matΔCxxC^* 2-cell embryos relative to respective *ctrl* samples. **E.** Barplot showing over-/under-representation and statistical significance of CGI-promoter genes belonging to the different DNAme clusters and being either significantly up- or down-regulated in *Kdm2a^KO^Kdm2b^KO^* relative to *ctrl* FGOs and/or in *Kdm2a^matKO^Kdm2b^matKO^* relative to *ctrl* 2-cell embryos for all, mat and pat specific sequencing reads. Numbers of significantly up- or down-regulated genes belonging to different DNAme clusters are indicated below the bars. Statistical significance is coded as follows: **** : p-val <= 0.001%; *** : p-val <= 0.01%; ** : p-val <= 0.1%; * : p-val <= 1%;. : p-val <= 5%. **F.** Barplot as in Figure 7E, for *Kdm2a^KO^Kdm2b^ΔCxxC^* FGOs and *Kdm2a^matKO^Kdm2b^matΔCxxC^* 2-cell embryos relative to respective *ctrl* samples. **G.** Genomic snapshots of *Gldc*, *Eomes*, *Wnt2* and *Ldlr* ^55^ as representative genes gaining DNAme at CGI promoters (highlighted in orange). RNA expression, DNA methylation and chromatin marks are indicated.

We next related expression changes of genes associated to these CGIs to aberrant DNAme. In FGOs, up- and down-regulated genes were rather evenly distributed among the different clusters (Figures 7C, 7D and S7H). As anticipated, aberrant DNAme was significantly associated with CGIs of clusters 1-4 genes that had been transcriptionally down-regulated in double mutant FGOs (Figures 7E and 7F). Thus, this likely reflects direct repression of CGIs functioning as promoters in oocytes by aberrant DNAme, gained through increased H3K36me2 in response to loss of H3K36me2 demethylase activities of KDM2A and KDM2B (Figure S7G). In single *Kdm2b^ΔCxxC^* FGOs, however, aberrant DNAme was not associated with transcriptional downregulation (Figure S7I), suggesting that KDM2A provides sufficient H3K36me2 demethylase activity to such single mutant GOs.

Counterintuitively, aberrant DNAme at other UCSC-annotated CGIs in clusters 2-5 was significantly associated with genes that were up-regulated in both double mutant FGOs and even in single *Kdm2b^ΔCxxC^* FGOs (Figures 7E, 7F and S7I). As described above for cluster 6 CGIs, the DNAme gain at such UCSC-annotated promoter CGIs probably reflects their localization within regions that become aberrantly transcribed in GOs from oocyte specific promoters located upstream of these UCSC-annotated CGIs and that are normally repressed by PRC1. Indeed, aberrant transcripts were elevated, not only downstream but also upstream of such CGIs (Figure S7G).

In two-cell embryos, genes associated with clusters 2-4 CGI promoters that had robust aberrant DNAme in *Kdm2a^KO^Kdm2b^KO^* and *Kdm2a^KO^Kdm2b^ΔCxxC^* oocytes were significantly down-regulated from maternal alleles but not from paternal alleles (Figures 7E and 7F). Contrasting to FGOs, aberrant DNAme was not associated with up-regulated expression in mutant two-cell embryos (Figures 7E and 7F). Hence, these results clearly indicate that aberrant maternal DNAme inherited from oocytes represses gene transcription during pre-implantation development. For example, zygotic expression of the glycine decarboxylase gene (*Gldc*), encoding a key enzyme in glycine metabolism^51–53^, is majorly suppressed in maternally deficient *Kdm2a*/*Kdm2b* two-cell embryos (Figure 7G). Other factors with well-known functions in blastocyst development (*Eomes*, *Gata4*, *Hnf1b*, *Junb*, *Klf4*, *Sox17*), placenta development (*Atrx*, *Hand1*, *Lhx3*, *Vash2*, *Wnt2*) and gastrulation (*Bmp4*, *Brachyury* (*T*), *Sox7*, *Tlx2*) are also hypermethylated at their promoters (Figures 7G, S7J and Table S7).

### Hypermethylation affects promoters of key developmental regulatory genes

Gene Ontology analysis of genes with hypermethylated CGI-promoters shows that many serve major functions throughout post-implantation embryonic development e.g., in cell fate specification and determination, morphogenesis, cellular differentiation and cell cycle regulation (Figure 7B and Table S6). They belong to various transcription factor gene families, such as *Foxo, Gata*, *Hand*, *Hes*, *Hox*, *Lhx*, *Nkx*, *Pax*, *Six*, *Sox, Tead*, *Tbx and Zfp*. Gene families involved in cell signaling, growth factor activity and cell adhesion were also affected (*Bmp*, *Dll, Fgf*, *Igf2*, *Pdgf*, *Pcdh*, *Rar*, *Wnt*, *Tgfb*), as were genes functioning in gonadal development and synaptonemal complex assembly. Hence, KDM2A and KDM2B protect large sets of CGI-promoters of key developmental regulators against hypermethylation in oocytes, thereby safeguarding development.

## DISCUSSION

In this study we identify the mechanism specifying the hypo-DNA-methylated genome characteristic of mouse oocytes. We further demonstrate the necessity of hypomethylation of the oocyte genome for embryonic development and correct gene expression following fertilization. Hypermethylation impairs zygotic gene expression and embryonic viability.

Our research reveals that in GOs global *de novo* DNA methylation acquisition beyond transcriptional units is in principle instructed by H3K36me2 (Figure 3). KDM2A and KDM2B serve critical roles in GOs in limiting H3K36me2 occupancy within and between genes as well as at gene promoters. Our data support that KDM2A/KDM2B demethylate H3K36me2 via their enzymatic activities^27,28^. In addition, by recruiting vPRC1.1 complexes that catalyze abundant and wide-spread H2AK119u1 on chromatin, KDM2A/KDM2B may also inhibit NSD and/or SETD2 histone methyltransferases from depositing H3K36me2/me3 ^42^.

At CGIs, we resolved the syntax of a multi-layered sequence and chromatin modifier interaction network specifying unmethylated versus methylated DNA states. In line with biochemical assays ^42,23^, our results show that KDM2A/KDM2B coordinate the balance between local H2AK119u1 and H3K36me2 levels, thereby controlling downstream *de novo* DNAme acquisition in growing oocytes, with local low H3K4me3 occupancy being permissive (Figure 4). Our work further shows that a selective set of CGIs are particularly vulnerable to aberrant H3K36me2 and DNAme acquisition. These CGIs are characterized by low CpGpG trinucleotide frequencies, which likely limit recruitment of the CFP1/SETD1B complex driving high H3K4me3 levels^43,39,45,42^.

It remains to be determined whether KDM2A and/or KDM2B serve related functions in male germ cells, thereby protecting CGIs also against erosion through deamination of methylated CpGs during evolution^2^. During gastrulation, *Kdm2b* and particularly *Mll2* were recently shown to partially suppress DNAme acquisition in the epiblast of gastrulating embryos^54^. While functional redundancy by paralogs needs to be considered, these findings point to a universal mechanism keeping H3K36me2 occupancy levels at CGIs low, thereby preventing aberrant CGI DNAme during the mammalian life cycle. Intriguingly, we observed aberrant DNAme at the CGI of the *Low-density lipoprotein receptor* (*Ldlr*) gene that was transcriptionally downregulated in *Kdm2a^KO^Kdm2b^KO^* (DNAme: 38.6%) and *Kdm2a^KO^Kdm2b^ΔCxxC^* (DNAme: 51.5%) FGOs compared to *ctrl* FGOs (DNAme: 0%) (Figure 7G and Tables S1-S3). Experimentally induced DNAme at the CGIs of the *Ldlr* and *Ankyrin repeat domain 26* (*Ankrd26*) genes was recently associated with transgenerational inheritance of reduced gene expression and metabolic phenotypes across multiple generations^55^. For *Ankrd26*, we measured 30% aberrant DNAme in *Kdm2a^KO^Kdm2b^KO^* FGOs (Table S3). At the *Ldlr* CGI, DNAme was readily detected in somatic tissues of transgenic animals, but was absent in primordial germ cells, oocytes, sperm and blastocyst embryos^55^. Instead, it became detectable in the epiblast of E6.5 gastrulation embryos^55^ arguing that at the *Ldlr* CGI an acquired epigenetic state other than DNAme confers epigenetic inheritance through the germline. Our work suggests that the presence of H3K36me2 and absence of H3K4me3 may constitute such memory.

Further, our study demonstrates the importance of the CXXC domain of KDM2B in recruiting vPRC1.1 to CGIs to deposit H2AK119u1. Interestingly, the absence of the CXXC domain did not prevent KDM2B^ΔCxxC^ from maintaining low H3K36me2 occupancy levels and precluding DNAme acquisition at genomic regions other than CGIs. Such activities likely reflect physiological functions of KDM2B beyond CGIs. Moreover, absence of KDM2A/KDM2B proteins provoked widespread de-repression of many PRC1/PRC2 target genes. Nonetheless, such transcriptional de-repression did not result in significant H3K36me3 deposition within gene bodies. As observed for the Set2 homologue in *Saccharomyces cerevisiae*^56^, SETD2 deposits H3K36me3 co-transcriptionally along with RNA polymerase 2, probably requiring multiple rounds of transcription to accumulate sufficient levels of the mark within gene bodies. In contrast, H3K36me2 was efficiently established within such lowly expressed genes as in intergenic regions, likely by one or more NSD family members.

Our study shows that KDM2A/KDM2B define the embryonic competence of oocytes required for proper pre-implantation development by restricting DNAme acquisition during oocyte growth. Given the lethality of maternal *Kdm2aKdm2b* mutants along the course of pre-implantation development, we suggest that the aberrantly gained maternal DNAme is maintained during this period, at least in a partially penetrant manner, reducing gene expression of key developmental regulators in a variegating manner between different embryos. In line with this, the extensive oocyte-derived DNAme is not majorly removed after fertilization in the zygote, unlike sperm-derived DNAme, suggesting that global DNAme reprogramming activities in early embryos can be restrained in a parent-of-origin specific manner (Figure 6A). Importantly, two-cell transcriptional profiling revealed a strong association between aberrant CGI promoter DNAme and suppression of maternal allele specific gene expression. The observed embryonic lethality implies haplo-insufficiency in autosomal gene expression in two-cell embryos, in which paternal expression is not sufficient to compensate for the loss of maternal expression caused by persistent promoter DNAme. This finding is in line with early lethality reported for mouse embryos with autosomal monosomy^57^. In addition, the aberrant promoter methylation observed in mutant oocytes at over 50 X-linked loci (including e.g., *Atrx*^58^) may effectively suppress expression in male and female early embryos and impair their development.

The developmental observations raise the intriguing question whether aberrant oocyte derived DNAme could in principle affect gene expression even in embryonic and/or placental tissues upon implantation. For example, reduced *WNT2* mRNA and protein expression, and increased promoter methylation in human placentas have been associated with early onset severe preeclampsia^59,60^. WNT2 expression is also reduced in villous tissues of patients with unexplained recurrent spontaneous abortions, which may impair trophoblast cell proliferation and migration via downregulating Wnt/β-catenin signaling^61^. In mice, deficiency for *Wnt2* causes placental defects^62^. Hence, for *WNT2* and possible other preeclampsia regulators, it will be important to investigate whether possible aberrant promoter DNAme measured in placental tissues originates from oocytes. CGI methylation in oocytes may then become diagnostic for their quality and embryonic competence.

## MATERIALS AND METHODS

### Mice

To generate maternally conditionally mutated mice, *Kdm2a^flox-JmJ^/^flox-JmJ^; Kdm2b^flox-JmJ^/^flox-JmJ^* mice and *Kdm2a^flox-JmJ^/^flox-JmJ^; Kdm2b^flox-CxxC^/^flox-CxxC^* (double-gene mutations) mice were crossed with mice carrying the Zp3-cre recombinase transgene, which excises floxed exons in growing oocytes^63^. *Kdm2a^flox-JmJ^/^flox-JmJ^; Kdm2b^flox-JmJ^/^flox-JmJ^; Zp3-cre* female mice produced oocytes deficient for KDM2A and KDM2B proteins. *Kdm2b^flox-CxxC^/^flox-CxxC^* mutation produces an in-frame *Kdm2b* transcript encoding a KDM2B protein lacking CxxC-motif binding domain. *Kdm2a^flox-JmJ^/^flox-JmJ^; Zp3-cre, Kdm2b^flox-JmJ^/^flox-JmJ^; Zp3-cre, Kdm2b^flox-CxxC^/^flox-CxxC^; Zp3-cre* (single-gene mutation) mice were segregated from double-floxed mice. Triple-gene mutations with *Dnmt1* or *Dnmt3a* were obtained by crossing double-floxed mice with *Dnmt1^flox^/^flox^* or *Dnmt3a^flox^/^flox^* mice, respectively^64,65^. All mutant mice were held on a C57BL/6J background.

We refer to *Ctrl* mice as genetically modified mice that we generated in experimental crosses but that do not harbor the *Zp3-cre* transgene. We refer to *wt* mice as genetically non-modified mice that we used in this study as sperm donors or that have been used in various published epigenomic studies as sources of gametes.

All experiments were performed in accordance with Swiss animal protection laws (licenses 2569, 2670, 3183, Gesundheitsdepartement Kanton Basel-Stadt, Veterinäramt, Switzerland) and institutional guidelines.

### Collection of mouse oocytes and preimplantation embryos

Growing oocytes (GOs) were collected from mice 9.0 or 14.0 days of age. Ovaries were dissociated in TrypLE Express Enzyme (1x) (Gibco; 12604013). Isolated oocytes were washed in M2 medium supplemented with Milrinone (25 μM, Sigma-Aldrich; 475840).

To collect fully grown germinal vesicle oocytes (FGOs) from 4- to 20-week-old female mice, 100 μl of Hyper Ova (CARD; KYD-010-EX-X5) or 5 I.U. of pregnant mare serum gonadotropin (PMSG, MSD; A207A01) were injected 46-52 h before the collection. Ovaries were dissected out in M2 medium (Merck; M7167) supplemented with Milrinone (25 μM) and cumulus cells were removed by mouth-pipetting using thin glass needles.

To collect MII oocytes, 7- to 20-week-old female mice were super-ovulated by injecting 5 I.U. of pregnant mare serum gonadotropin and 5 I.U. of human chorionic gonadotropin (hCG, MSD; A201A01). For the examination of preimplantation embryos, we performed *in vitro* fertilization (IVF) to synchronize fertilization timing across experimental conditions. To generate hybrid strain embryos, JF1/MsJ (The Jackson Laboratory; 003720) strain males were used. Spermatozoa were capacitated in Human Tubal Fluid medium (HTF) (Merck Millipore; MR-070-D) supplemented with 10 mg/ml Albumin (Sigma; A-3311) for 1-1.5 h prior to insemination and used for IVF performed in HTF with Albumin. The starting time of insemination was designated as 0 hpf (hours post-fertilization). At 4 hpf, eggs were transferred to KSOM medium (Millipore; MR-106-D) and the number of ovulated MII oocytes was counted. The formation of two pronuclei was visually confirmed under the microscopy at 6 hpf. Fertilized eggs were cultured in KSOM medium covered with mineral oil (Sigma; M5310) at 37°C with 5% CO^2^ and 5% O^2^ air. Preimplantation embryo development was observed at embryonic developmental days e1.25, e2.25, e3.25, e4.25 and e5.0 after IVF. Only for collecting *Kdm2a^flox-JmJ^/^flox-JmJ^; Kdm2b^flox-CxxC^/^flox-CxxC^* maternally deficient 2-cell embryos for smartseq2 RNA sequencing, embryos were generated by Intra cytoplasmic sperm injection (ICSI) using JF1/MsJ spermatozoa to prevent the contamination of RNA from multiple sperm strongly attached to the blastomere surface, which can occur after IVF.

To collect samples for genomics and immunostaining experiments, oocytes and preimplantation embryos were first treated with acidic Tyrode’s solution (Sigma-Aldrich; T1788) supplemented with 0.01% polyvinyl alcohol (Sigma-Aldrich; P8136) to remove the zona pellucida, and then washed in M2 medium.

For collecting samples for smartseq2 and WBGS, the zona pellucida was removed as described. FGOs and late two-cell embryos (at 30 hpf) were subsequently washed in M2 medium and PBS supplemented with 0.01% PVA. This was followed by cell lysis in smartseq2 lysis buffer or RLT plus buffer (QIAGEN), respectively.

### Immunofluorescence staining of oocytes and preimplantation embryos

After removing the zona pellucida, collected GOs, FGOs and embryos were washed in PBS supplemented with 0.05% PVA (PBS-PVA). Fixation was done in 4% paraformaldehyde (PFA) in PBS-PVA at room temperature for 15 min. After washing in PBS-PVA three times, samples were permeabilized in 0.5% Triton X-100/ PBS at room temperature for 15 min followed by washing in PBS containing 0.1% Tween-20 (Sigma-Aldrich; P2287) (PBS-T). For the staining of 5mC, samples were treated with 4N HCl for 10 min followed by incubation in 100mM Tris-HCl (pH 8.0) for 10 min, both at room temperature. 2% BSA (w/v) or 5% normal goat serum in PBS-T was used for blocking and the incubation with primary antibodies diluted in PBS-T with 1% BSA or 5% normal goat serum was done at 4 °C overnight. The following primary antibodies were used: anti-H2AK119ub1 (1:20,000; Cell Signaling Technology; 8240), anti-H3K27me3 (1:15,000; Cell Signaling Technology; 9733), anti-Kdm2a (1:500; Abcam; ab191387), anti-H3K36me2 (1:500; MBL International; MABI0332), anti-H3K36me3(1:1,000; Abcam; ab9050), anti-5mC (1:500; Eurogentec; BI-MECY-100). After washing out the primary antibodies in PBS-T, the secondary antibody incubation was performed in a lightproof box at room temperature for 2 h.

Secondary antibodies (Thermo Fischer Scientific) were Alexa Fluor (AF) 488 donkey anti-mouse IgG, AF488 donkey anti-rabbit IgG, AF568 donkey anti-mouse IgG, AF568 donkey anti-rabbit IgG, AF647 donkey anti-mouse IgG and AF647 donkey anti-rabbit IgG at 1:1,000 dilution in PBS-T. After washing three times in PBS-T, samples were mounted on a glass slide in Vectashield Antifade Mounting Medium with DAPI (Vector Laboratories; H-1200, H-2000). Z-stack fluorescent images (0.33 µm steps) were acquired by Axio Imager M2 (ZEISS) combined with a Yokogawa CSU W1 Dual camera T2 spinning disk confocal scanning unit (YOKOGAWA). Projection image processing, signal intensity and area size quantifications were done using Fiji software. The signal intensity within nuclei of each Z-stack plane was calculated and normalized by the DAPI positive area size. In box plots, whiskers extend to data points that are less than 1.5 x interquartile range away from the 1st/3rd quartile.

### Smart-seq2 RNA-sequencing of growing and fully grown oocytes and two-cell embryos

FGOs and genetically hybrid two-cell embryos were prepared as described above. Libraries were prepared following the Smartseq2 protocol^66^. For each genotype, we prepared 20-25 libraries, each from one individual FGO or embryo. After the removal of the zona pellucida as described above, samples were washed in PBS (Lonza; 11629980) supplemented with 0.01% PVA (PBS-PVA) once and lysed in 4ul of lysis buffer composed of 0.09% Triton-X 100 (Sigma-Aldrich; T9284), 2U SUPERase IN RNase inhibitor (Invitrogen AM2694), 2.5μM Oligo-dT primer (5’-AAG CAG TGG TAT CAA CGC AGA GTA CTT TTT TTT TTT TTT TTT TTT TTT TTT TTT TVN; Microsynth AG), dNTP mix (2.5mM each, Promega; U1515), ERCC RNA Spike-In Mix (1:3.2x10^7^ Thermo Fischer Scientific 4456740) in 8-well strips or 96-well plate (one sample per one well). Samples were immediately frozen on dry ice and kept in the -80C freezer for long-term storage. After thawing, lysed samples were denatured at 72°C for 3 min and quickly chilled on ice. Reverse transcription mix composed of 100U SuperScript II reverse transcriptase (Thermo Fischer Scientific, 18064014), 5U SUPERase IN RNase inhibitor, 1X Superscript II first-strand buffer, 5mM DTT (in SuperScript II reverse transcriptase), 1M Betaine (Sigma, B0300-1VL), 6mM MgCl2, 1μM template-switching oligos (TSOs) (5’AGCAGTGGTATCAACGCAGAGTACATrGrG+G–3’; EXIQON) was added to samples to obtain a total volume 10μl and the reverse transcription was performed in PCR machine. Next, PCR pre-amplification was performed by adding 1X KAPA HiFi HotStart Ready Mix (KAPA Biosystems, KK2602), 0.1μM ISPCR primers (5’-AAGCAGTGGTATCAACGCAGAGT; Mycrosynth AG) in total volume 25μl. The preamplification PCR cycle numbers were 14-16 cycles for FGOs and 16 cycles for two-cell embryos. Half of amplified DNA was purified with SPRI AMPure XP beads (sample to beads ratio 1:1, Beckman; A63881) and eluted in 15μl Buffer EB (QIAGEN). 1ng of pre-amplified DNA was used for tagmentation reaction (7min @55°C) using Tn5 tagmention mix (1x TAPS-DMF buffer, self-purified Tn5-tagmentase (1:1,200)) in total volume 20μl. The reaction was stopped by adding 5μl of 0.2% SDS and kept at 25°C for 7 min. Adapter-ligated fragment amplification was done using Nextera XT index kit v2(Illumina) in a total volume 50μl (1x Phusion HF Buffer, 2U of Phusion High Fidelity DNA Polymerase (Thermo Fischer Scientific; F530L), dNTP mix (0.3mM each)) with 9-10 cycles of PCR. Library DNA was purified by SPRI AMPure XP beads (sample to beads ratio 1:1) and eluted in 12μl Buffer EB. Sequencing was performed on an Illumina HiSeq2500 machine with single-end 75bp read length or NovaSeq6000 machine with paired-end 2x50bp read length (Illumina).

### Total RNA sequencing of growing oocytes

GOs were isolated from the d14.0 mice. 60-80 oocytes were pooled and frozen in 100μl of Buffer RL. 4 and 3 biological replicates were prepared for *ctrl* and *Kdm2a^KO^Kdm2b^KO^*, respectively. RNA was purified by using Single Cell RNA Purification Kit (NORGEN; 51800). Libraries were prepared according to Illumina Stranded total RNA-seq protocol with Illumina IDT DNA/RNA UDI indexes. Sequencing was performed on NovaSeq6000 machine with paired-end 2x50bp read length (Illumina).

### CUT&RUN of oocytes

CUT&RUN libraries were prepared as previously described^67^. 300 to 500 FGOs were used for H2AK119u1 and 200 FGOs were used for H3K36me2 and H3K36me3 for each Cut and Run library. For each genotype and histone PTM, at least two independent libraries were prepared. The antibodies were rabbit anti-H2AK119ub1 (1:100; Cell Signaling Technology; 8240), mouse anti-H3K36me2 antibody (1:100; Cosmo bio; MABI0332) and anti-H3K36me3(1:100; Abcam; ab9050). Self-purified Protein AG-MNase (pDNA was from Addgene; 123461) was used. CUT&RUN libraries were prepared using NEBNext Ultra II DNA Library Prep Kit for Illumina (NEB; E7645L) and sequenced on NextSeq500 (paired-end, 2x75 bp) or NovaSeq6000 (paired-end 2x50bp).

### Whole Genome Bisulfite sequencing (WGBS) of oocytes

FGOs were prepared as described above. WGBS was performed as described previously^68^ with modifications. For bisulfite conversion, EZ DNA Methylation-Direct Kit (Zymo Research; D5020) was used. After zona pellucida removal, we generated 13 pools of 10 FGOs per genotype in a single well of 8-well strips containing 2.5μl of Buffer RLT (QIAGEN; 79216). Samples were stored in a -80C freezer. After thawing, 7.5μl of Nuclease Free Water (Invitrogen; AM9937) and 65μl of CT conversion reagent (EZ DNA Methylation-Direct Kit; Zymo Research D5020) were added and samples were incubated in a PCR machine (98°C 8 min, 65°C 180 min). DNA was purified using PureLink micro kit (Thermo Fischer Scientific; K310050). DNA bound to the purification columns was washed by wash buffe**r (**PureLink micro kit) once, 100μl of M-Desulphonation buffer (EZ DNA-Methylation kit) was applied on the columns and the incubation was done at room temperature for 15 min to complete the CT-conversion. DNA on the columns was further washed twice and then eluted using the DNA strand synthesis mix (4μl of Blue Buffer (Enzymatics), 1.6μl of dNTP (10mM each) (Roche; 4638956001), 1.6μl of 20μM Preamp primer (5’-[Btn]TGACTGGAGTTCAGACGTGTGCTCTTCCGATCTNNNNN*N, SIGMA) and 32.8μl of Nuclease free water). DNA mixture was denatured at 65°C for 3min and quickly chilled. Then 1μl of Klenow Fragment (3’-5’ exo-) (Enzymatics; P7010-LC-L) was added and the strand synthesis reaction was done in the PCR machine with the following program (4°C for 5min, 4°C rising to 37°C with increasing 1°C every 15 sec, 37°C for 30 min, 4°C pause). Another 4 rounds of the strand synthesis reaction were performed, in that synthesized DNA from the previous PCR cycle was denatured at 95°C for 1 min and quickly chilled, then added 2.4μl of master mix (0.25μl of 10x Blue Buffer, 0.1μl of dNTP, 1μl of 20μM Preamp primer, 0.4μl of Klenow Fragment (3’-5’ exo-) and 0.65μl of water) before the strand synthesis PCR. Strand synthesized DNA was treated with 40U of Exonuclease I (NEB; M0293S). DNA was purified by SPRI AMPure XP beads (sample to beads ratio 0.8:1) and eluted by the 2^nd^ strand synthesis mix (5μl of 10x Blue Buffer, 2μl of dNTPs (10mM each), 2μl of 20μM Adapter primer 2 (5’-ACACTCTTTCCCTACACGACGCTCTTCCGATCTNNNNN*N, SIGMA) and 39μl of water). DNA was denatured at 95°C for 45sec and quickly chilled. Then 2μl of Klenow fragment was added and amplification incubation was done with the same program with the first strand synthesis. KAPA HiFi HotStart PCR Kit (Roche; KK2502) was used for indexing-amplification with NEBNext Multiplex Oligos for Illumina (NEB) by 10-12 PCR cycles. Libraries were purified by SPRI AMPure XP beads (sample to beads ratio 0.8:1) and eluted in 12μl Buffer EB. WGBS libraries were sequenced on NextSeq with single-end 75bp read length (Illumina).

### Alignment and quantification of RNA-Seq data of oocyte samples

RNA-Seq datasets were aligned to the Mus musculus genome assembly (GRCm38/mm10 Dec. 2011) as single-end (Smart-Seq2 polyA datasets, Figures S2A-S2B, S5B, S7A-S7dD) or paired-end (total random primed datasets, Figure 5C) using STAR^69^ with parameters “-outFilterMultimapNmax 300 - outMultimapperOrder Random -outSAMmultNmax 1 -alignIntronMin 20 -alignIntronMax 1000000”. Expression quantification for genes in Bioconductor annotation package TxDb.Mmusculus.UCSC.mm10.knownGene (version 3.2.2) was done using QuasR R package (Gaidatzis et al. 2015) selecting only uniquely mapped reads (mapqMin=255). RPKM values for genes were calculated by normalizing exonic read counts to total exonic length of each gene and total number of reads mapping to all exonic regions in each library. RPKM values were log2 transformed using formula log2(RPKM + psc) – log2(psc) where pseudo-count psc was set to 0.1.

### Alignment and allelic assignment for RNA-Seq data of two-cell embryo samples

Smart-seq2 RNA-Seq samples for hybrid Bl6 x JF1 F1 2-cell embryos were separately aligned to Bl6 and JF1 genomes obtained by incorporating JF1 single-nucleotide polymorphisms (SNPs) into reference mm10 genome using previously published SNP table^70^ RNA-seq reads were categorized as maternal (Bl6), paternal (JF1) or undefined based on minimal number of mismatches in alignments to both genomes. Total number of maternal and paternal reads was used as library size for calculating RPKM values and differential expression analysis.

### Differential expression analysis of RNA-seq datasets for single FGOs

Genes with at least 1 read per million in at least 3 samples were included in the statistical analysis of differential expression. edgeR^71^ was used for statistical analysis of differential gene expression between *Kdm2a^KO^Kdm2b^KO^*, *Kdm2a^KO^Kdm2b^ΔCxxC^*, *Kdm2b^ΔCxxC^* and respective *ctrl* FGOs. Generalized Linear Model was fit using genotypes as covariates. Statistical significance was estimated using log-likelihood tests and the Benjamini-Hochberg method was used to correct for multiple testing.

### Differential expression analysis of RNA-seq datasets for single day 9 and day 14 GOs

Principal component analysis for transcriptomes of single day 9 and day 14 GOs revealed strong dependence on oocyte diameter which explains 16% of variance (Figure S5B). To take this variation into account we included basis functions for natural splines into a generalized linear model. Analysis for day 9 and day 14 growing oocytes was performed separately, and model matrix was constructed by 1) including oocyte genotypes and 2) including basis functions for natural splices with 3 components generated by ns function in R package splines (version 3.5.1) using oocyte diameters as knots. More explicitly, design matrix for GLM was generated using model.matrix function with formula ∼ 0 + genotype + ns(d,3), where d is a diameter of each single oocyte. To control possible overfitting, we performed the same analysis using randomly shuffled oocyte diameters.

### Differential expression analysis RNA-seq datasets for 2-cell embryos

To consider possible developmental delays due to random experimental factors or indirect effects of maternal depletion of *Kdm2a* and *Kdm2b* we additionally profiled gene expression in *ctrl* single embryos at early and late 2-cell stages (Figures S7A-S7D), which served as a timing control and allowed us to study effects of maternal genetic mutation of *Kdm2a* and *Kdm2b* in the context of gene expression changes which normally occur during maternal-to-zygotic transition in 2-cell embryos. Pseudotime for each embryo was estimated using R package SCORPIUS (v1.0.8)^72^ using read counts for both exonic and intronic regions of genes and removing genes in chrX and chrY as well as imprinted genes annotated in geneimprint website (https://geneimprint.com/site/genes-by-species.Mus+musculus).

To consider possible effects of embryo sexes on gene expression we identified sex of each embryo using proportions of reads mapping to chrX and chrY.

Genes with at least 1 read per million in at least 3 samples were included in the statistical analysis of differential expression which was done using edgeR package^71^. Construction of a model matrix for generalized linear model (GLM) was done by 1) including interaction between genotypes and embryo sexes as covariates and 2) including basis functions for natural splines with 3 components generated by ns function in R package splines (version 3.5.1) using pseudotime as knots to regress out effects of possible developmental delays. More explicitly, design matrix for GLM was generated using model.matrix function with formula ∼ 0 + genotype:sex + ns(PsT,3). To control possible overfitting by splines, the same model was fit for samples with randomly shuffled pseudotime estimates.

Expression changes (log2(Fold-changes)) and FDR were calculated for difference between sex averaged coefficients for maternal *mutants* and respective *ctrls* using log-likelihood test and Benjamini-Hochberg method for multiple testing correction.

### Gene Ontology enrichment analysis

Enrichment analysis for Gene Ontology terms was done using R package topGO (version 2.48.0)^73^ with parameters method=”weight01” and statistic=”fisher” extracting Gene Ontology gene annotation from the Bioconductor Annotation Package org.Mm.eg.db (version 3.15.0)^74^ (Table S7) or mapping to slim Gene Ontology using map2slim tool (Figures 7B, S3A, Table S6).

Visualization of Gene Ontology enrichments was done by calculation of pairwise Jaccard distances between significant Gene Ontology terms based on intersections and unions of significantly affected gene sets having corresponding Gene Ontology term annotations. After pairwise Jaccard distances between Gene Ontology terms were calculated we applied multidimensional scaling (MDS) using R function cmdscale and represented Gene Ontology terms on a 2D plot where size was scaled by obs./exp. ratio, color was chosen to reflect statistical significance and relative position reflects similarities in gene sets (Figures 7B, S3A).

### Alignment and quality control of WGBS datasets

The quality of the data was assessed using FastQC (v0.11.8) and adapters were trimmed using TrimGalore (v0.6.2)^75^ with parameters “--illumina --stringency 5 --clip_r1 6 --three_prime_clip_r1 6”. Alignment to mm10 genome and deduplication was done using Bismark (v0.22.3) ^76^ with parameters “--local --non_directional” (Figure S4B). Reproducibility of samples has been assessed by calculating levels of DNAme in 1e+5 randomly selected 500bp genomic tiles, calculating pairwise Euclidean distances between samples and performing multidimensional scaling for obtained distance matrix (Figure S4A). Finally, samples with small library sizes (≤ 10e+6 reads), insufficient bisulfite conversion efficiency estimated using total levels of non-CpG (CHG, CHH) methylation (≥7%) as well as outlier samples were removed from the analysis and remaining samples for each genotype were merged for further analysis.

### Analysis of CUT&RUN sequencing data

Reads from the CUT&RUN experiments were aligned to a composite mouse-fly genome (GRCm38/mm10 and dm6 UCSC assemblies, https://genome.ucsc.edu) using the qAlign function of the QuasR R-bioconductor package with the “paired” parameter set to ‘fr’ and the remaining parameters set to defaults.

To account for coverage distortions across the different CUT&RUN libraries we employed a procedure whereby counts where normalized to the total number of reads originating from regions (gene body + 5kbp flanking windows) of genes stably expressed across genotypes according to the matching RNAseq data (absolute Log2CPM > 0.25, absolute Log2FoldChange < 0.25). This procedure assumes that chromatin marks over stably expressed regions remain -on average-unchanged. In the case of counts over genomic tiles normalized counts were also min-shifted to the 15th centile to account for detection-limit and background signal level differences across libraries.

For the presented results libraries of biological replicates of either wild-type or mutant libraries have been merged unless otherwise specified.

### Genome arithmetic operations

Read counting over specified genomic intervals, or genomic tiles of the mouse genome (GRCm38/mm10) was carried out with the QuasR count function with the “orientation” parameter set to ‘any’ and default parameters otherwise, excluding non-canonical chromosomes.

Gene and CpG-island coordinates for overlap counting and other genome-arithmetic operations were taken from the UCSC annotation database (mm10.knownGene and mm10 CpG Island track respectively, https://genome.ucsc.edu), unless otherwise specified. Specifically, for CpG-island counting operations the regions were resized to windows with length equal to the median CpG island width (533nt) preserving their center coordinate.

### Clustering and heatmaps for chromatin, sequence and expression features

Clusters presented on the different subsets of genes / intergenic regions were determined using k-means clustering with 100 random starts using the kmeans implementation of R stats on standardized features. Heatmaps of genomic tiles were plotted using the Heatmap function of the ComplexHeatmap R-Bioconductor package. Plotted tiles were smoothed with a running mean smoothing kernel of width 5. Color-scales were thresholded on both low and high values at the 2nd and 98th centile respectively.

### Regularized linear regression for chromatin-mark and DNA-methylation-modelling

For the modelling of basal methylation levels, differential methylation levels and differential H3K36me2 levels across genotypes we opted for the lasso regression framework to identify predictive explanatory variables. Since multicollinearity was extensive among the set of predictors, we also describe for each prediction task the covariance structure of the independent variables to facilitate model interpretability. For all models, CpG island chromatin and sequence features were calculated on the 533nt resized set of CpG islands except for the lower coverage CUT&RUN datasets (H3K36me2, H2AK119ub1) where the signal was calculated on a larger window (1066nt) to account for the lower resolution. For gene-body (gb) and RNA-seq expression features the signal refers to gene-bodies of the nearest annotated genes. All independent variables were z-score normalized prior to fitting.

For the lasso regression fitting we used the CRAN glmnet R package implementation. Briefly, in a first step we selected the lambda parameter as the largest value of lambda such that error is within 1 standard error of the minimum (lambda.1se) in a 10-fold cross validation. We next performed lasso regression with the selected lambda parameter.

### Example commands

*CV_fit <-cv.glmnet(X, Y, alpha=1, nfolds=10, lambda= 10^seq(-2, -6, by = -.05)) lambda_best <-CV_fit$lambda.1se*

fit <-glmnet(x, y, lambda=lambda_best)

In the case of modelling differential methylation or differential H3K36me2 levels we used a setting where the response variable is the mutant genotype levels while including wildtype levels as a predictor. This choice circumvents the issue of selecting features that are predictive of wild-type levels of the dependent variable (as opposed to differential levels) which is inadvertently the case when one models directly differential levels in the absence of the wild-type levels among the set of predictors. One alternative implementation -that yields almost identical results to the ones presented here (data not shown)-is to first regress out the effect of wild-type levels on mutant signal levels and subsequently model the residual on the remaining set of predictors. In our implementation we omit the wild-type signal levels when we present the most important predictors.

### Data visualization

Chromatin, sequence and expression feature heatmaps were generated with the ComplexHeatmap R bioconductor package. Chromatin features are Z-score transformed except when otherwise indicated. In heatmaps in Figures 2I, 3F, S3D and S4E, we included only genes with gene bodies > 5Kbp and < 80kbp in order to reduce plotting artefacts. Calculations were performed on 10 equal width (500bp) windows for both the regions of 5kbps upstream of the TSS and 5kbps downstream of the TTS. Calculations were performed on 20 variable width windows for the full gene-body regions. Prior to plotting local smoothing of the genomic signals was performed using a local mean smoother with a kernel size of 5 windows.

Boxplots were generated with the ggplot2 *geom_boxplot* function. Whiskers extend to 1.5 the IQR range.

### Chromatin and sequence analysis at UCSC-annotated CGIs

Enrichments for H2AK119u1 in *wt* FGO, H3K4me3 in *wt* GO as well as H3K36me2 in *ctrl* and *mutant* FGO were calculated in 533bp regions around the center of 16’023 CGIs. CGIs were split into groups with low (7,506 CGIs) and high (8,517 CGIs) H2AK199u1 levels based on calculated enrichments in *wt* FGO. Next, CGIs were split into 10 groups according to H3K4me3 levels in *wt* GO such that each group contains similar number of CGIs (1,439 – 1,842) using function cut_number from ggplot2 R package. Finally, for each H3K4me3 group separately for low and high H2AK119u1 groups we plotted boxplots for H3K36me2 in *ctrl* and *mutant* FGOs as well counts of CCG and CGG trinucleotides normalized per 100bp (Figure 5D).

### Quantification and statistical analysis

Statistical analyses were performed using R. All statistical tests, p values and sample numbers were stated in figure panels or legends. Statistical p values were calculated using two-tailed Student’s t-test and Tukey’s HSD test in immunostaining signal intensity comparison. Fisher’s exact test was used for the comparisons of embryonic development results.

## Data and code availability

All 597 genomic data sets (RNA-seq, WGBS and CUT&RUN data samples) are available at GEO under GSE234968 (https://www.ncbi.nlm.nih.gov/geo/query/acc.cgi?acc=GSE234968).

## Supporting information

Kawamura_Peters_Kdm2a-Kdm2b_Supplemental data

## ACKNOWLEDGMENTS

We thank B. Knowles, E. Li and R. Jaenisch for providing *Zp3-cre* transgenic mice, *Dnmt3a* and *Dnmt1* conditionally deficient mice, respectively. We gratefully acknowledge A. Inoue for sharing the CUT&RUN protocol and S. Henikoff for providing for the pAG/Mnase plasmid (Addgene 123461). We thank S. Bourke, J. Eglinger and L. Gelman (Facility for Advanced Imaging and Microscopy) and members of the FMI animal facility for excellent assistance. We thank M. Bühler, D. Schübeler, P. de Boer and laboratory members for critical reading of the manuscript. This research was supported by Japan Society for the Promotion of Science fellowship (Y.K.K), Naito fellowship (Y.K.K), Novartis Research Foundation, the Swiss National Science Foundation (406340_128131) and the European Research Council (ERC) under the European Union’s Horizon 2020 research and innovation programme (grant agreement ERC-AdG 695288 – Totipotency).

## AUTHOR CONTRIBUTIONS

Y.K.K. and A.H.F.M.P. conceived the study. Y.K.K., E.A.O., P.P. and A.H.F.M.P. designed the experiments and interpreted the data. Y.K.K. performed genetic, cell biology and genomic experiments. P.P and E.A.O. performed computational data analyses. T.K. and H.K. provided *Kdm2a* and *Kdm2b* conditional mutant strains. N.N. purified protein-AG-MNase. M.B.S. supported computational analyses. S.A.S. assisted and supervised genomic sequencing. A.H.F.M.P. supervised the project and wrote the manuscript with input from all authors.

## INCLUSION AND DIVERSITY

We support inclusive, diverse and equitable conduct of research.

## RESOURCES AND REAGENTS

Further information and requests for resources and reagents should be directed to Antoine Peters

