## Supplementary material for "Preventing CpG island hypermethylation in oocytes safeguards mouse development": Kawamura_Peters_Kdm2a-Kdm2b_Supplemental data

#### Supplementary Figures and Supplementary Table Legends

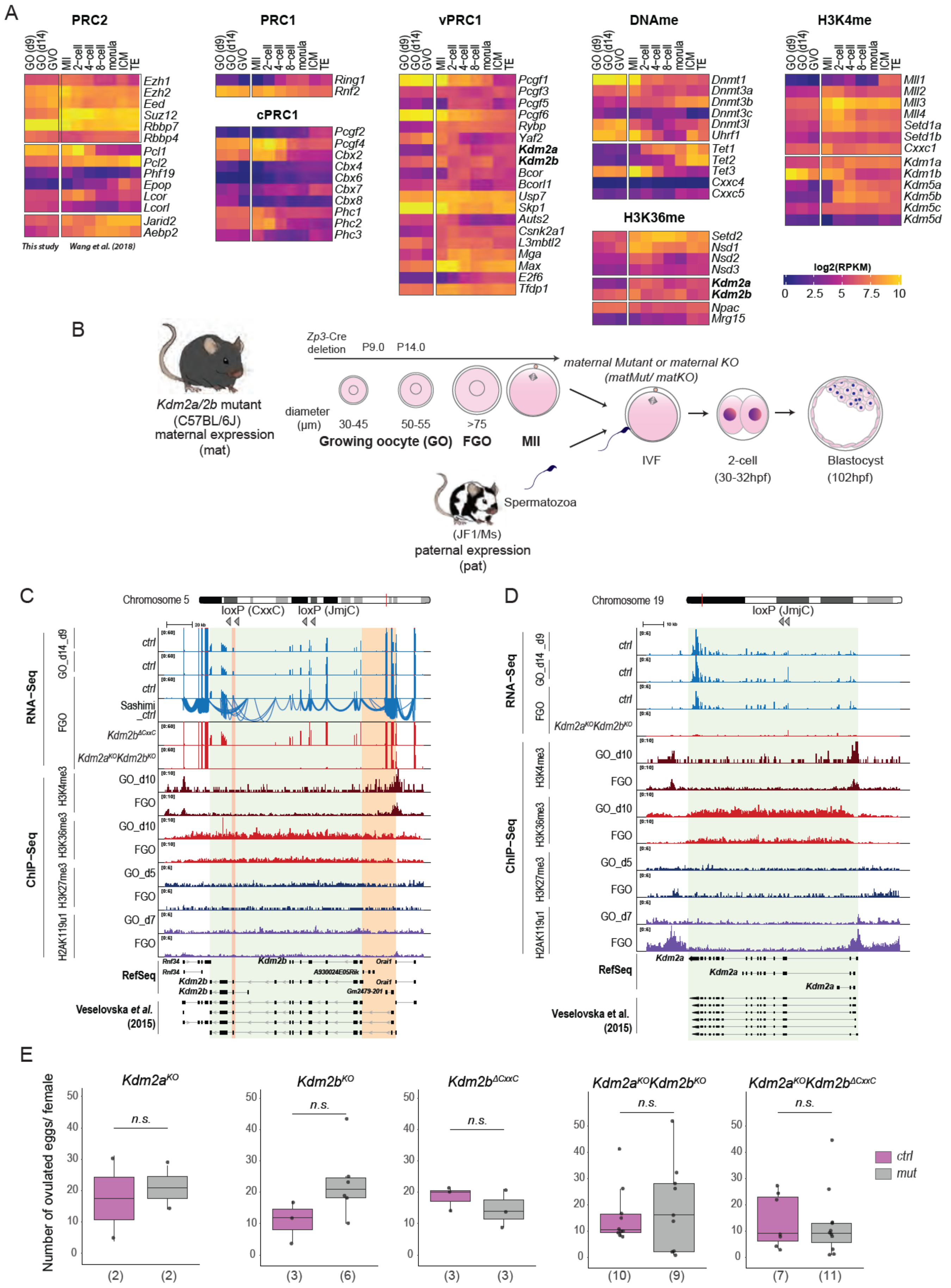
A
PRC2
PRC1
cPRC1
vPRC1
DNAme
H3K36me
H3K4me
GO (d9)
GO (d14)
GVO
MII
2-cell
4-cell
8-cell
morula
ICM
TE
Ezh1
Ezh2
Eed
Suz12
Rbbp7
Rbbp4
Pcl1
Pcl2
Phf19
Epop
Lcor
Lcorl
Jarid2
Aebp2
This study
Wang et al. (2018)
Ring1
Rnf2
Pcgf2
Pcgf4
Cbx2
Cbx4
Cbx6
Cbx7
Cbx8
Phc1
Phc2
Phc3
Pcgf1
Pcgf3
Pcgf5
Pcgf6
Rybp
Yaf2
Kdm2a
Kdm2b
Bcor
Bcorl1
Usp7
Skp1
Auts2
Csnk2a1
L3mbtl2
Mga
Max
E2f6
Tfdp1
Dnmt1
Dnmt3a
Dnmt3b
Dnmt3c
Dnmt3l
Uhrf1
Tet1
Tet2
Tet3
Cxxc4
Cxxc5
Setd2
Nsd1
Nsd2
Nsd3
Kdm2a
Kdm2b
Npac
Mrg15
Mll1
Mll2
Mll3
Mll4
Setd1a
Setd1b
Cxxc1
Kdm1a
Kdm1b
Kdm5a
Kdm5b
Kdm5c
Kdm5d
log2(RPKM)
0 2.5 5 7.5 10
B
Kdm2a/2b mutant
(C57BL/6J)
maternal expression
(mat)
Zp3-Cre deletion
P9.0 P14.0
diameter (µm)
30-45
50-55
>75
Growing oocyte (GO)
FGO
MII
maternal Mutant or maternal KO
(matMut/ matKO)
IVF
2-cell
(30-32hpf)
Blastocyst
(102hpf)
Spermatozoa
(JF1/Ms)
paternal expression
(pat)
C
Chromosome 5
loxP (CxxC)
loxP (JmjC)
RNA-Seq
ChIP-Seq
GO_d14_d9
FGO
ctrl
Sashimi_ctrl
Kdm2bΔCxxC
Kdm2aKOKdm2bKO
H3K4me3
H3K36me3
H3K27me3
H2AK119u1
GO_d10
GO_d5
GO_d7
RefSeq
Rnf34
Kdm2b
Orai1
A930024E05Rik
Gm2479-201
Veselovska et al. (2015)
D
Chromosome 19
loxP (JmjC)
Kdm2a
E
Kdm2aKO
Kdm2bKO
Kdm2bΔCxxC
Kdm2aKOKdm2bKO
Kdm2aKOKdm2bΔCxxC
Number of ovulated eggs/ female
n.s.
(2) (2)
(3) (6)
(3) (3)
(10) (9)
(7) (11)
ctrl
mut

**Figure S1: Deficiency of *Kdm2a* and/or *Kdm2b* does not impair oogenesis, related to Figure 1.**

- **A.** RNA expression of multiple genes in GOs (at day 9, day 14), FGOs and pre-implantation embryos at different stages of development [77].
- **B.** Breeding scheme to generate embryos maternally deficient (*matKO*) for protein function by crossing conditional $Kdm2a^{KO}$, $Kdm2b^{KO}$, $Kdm2b^{\Delta CxxC}$, double or compound mutant females (all on C57BL/6J genetic background) with *wt* males (on a C57BL/6J or JF1 genetic background).
- **C.** RNA expression and chromatin status along the RefSeq-annotated *Kdm2b* locus (in reverse orientation; highlighted in green) in *ctrl* and various *Kdm2b* conditionally mutant oocytes. *Kdm2b* and neighboring *Rnf34* and *Orai1* genes are highly expressed throughout oogenesis and are associated with high H3K4me3 occupancy at gene promoters, widespread H3K36me3 enrichment along gene bodies and absence of repressive H3K27me3 and H2AK119u1. Chromatin and splice-junction analysis revealed that *Kdm2b* transcription in oocytes initiates from an alternative promoter, encoding a protein that is 54 amino acids longer than the canonical form[31,40]. The promoter also drives the expression of the short *Gm2479-201* transcript. The region encompassing the alternative transcriptional start site (TSS) as well as the exon encoding the CXXC domain of *Kdm2b* are highlighted in orange. The positions of LoxP sites in the floxed *JmjC* (referred to as *KO*) and *CxxC* conditional alleles are indicated.
- **D.** RNA expression and chromatin status along the RefSeq-annotated *Kdm2a* locus (in reverse orientation; highlighted in green) in *ctrl* and *Kdm2a* conditionally deficient oocytes. *Kdm2a* is expressed throughout oogenesis and is marked by H3K4me3 at its promoter and by H3K36me3 along its gene body. While H3K27me3 is absent, H2AK119u1 levels are increased upon oocyte growth in FGOs.
- **E.** Ovulation rates of single and double conditionally mutant females upon hormonal superovulation treatment. Numbers of analyzed females are indicated. P-values according to two-sided student's *t*-test.

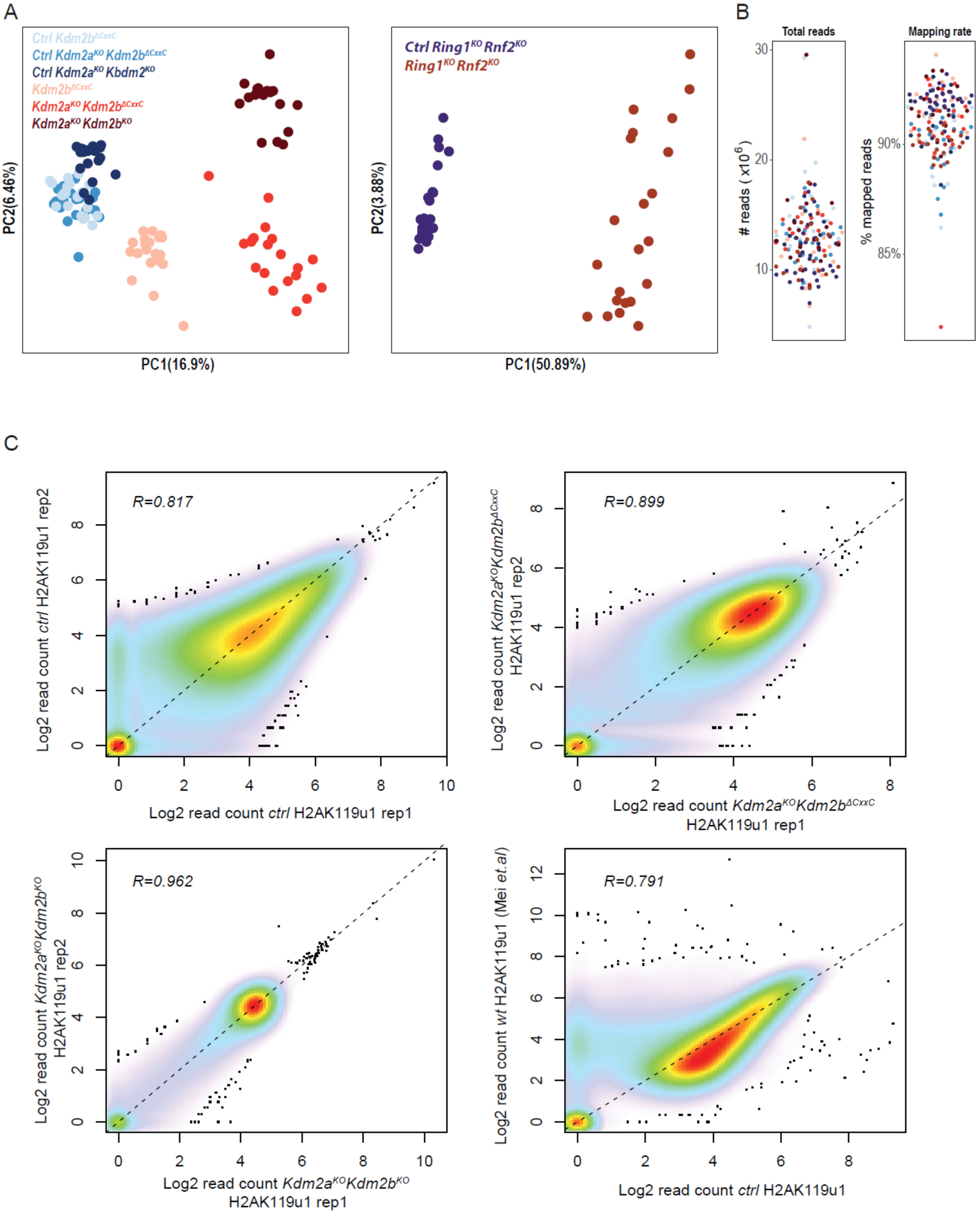
A
Ctrl Kdm2b^ΔCxxC
Ctrl Kdm2a^KO Kdm2b^ΔCxxC
Ctrl Kdm2a^KO Kbdm2^KO
Kdm2b^ΔCxxC
Kdm2a^KO Kdm2b^ΔCxxC
Kdm2a^KO Kdm2b^KO
PC2(6.46%)
PC1(16.9%)
Ctrl Ring1^KO Rnf2^KO
Ring1^KO Rnf2^KO
PC2(3.88%)
PC1(50.89%)
B
Total reads
### reads ( x10^6 )
30
20
10
Mapping rate
% mapped reads
90%
85%
C
R=0.817
Log2 read count ctrl H2AK119u1 rep2
Log2 read count ctrl H2AK119u1 rep1
R=0.899
Log2 read count Kdm2a^KO Kdm2b^ΔCxxC H2AK119u1 rep2
Log2 read count Kdm2a^KO Kdm2b^ΔCxxC H2AK119u1 rep1
R=0.962
Log2 read count Kdm2a^KO Kdm2b^KO H2AK119u1 rep2
Log2 read count Kdm2a^KO Kdm2b^KO H2AK119u1 rep1
R=0.791
Log2 read count wt H2AK119u1 (Mei et.al)
Log2 read count ctrl H2AK119u1

**Figure S2: Quality control analyses of RNA-seq and CUT&RUN-seq data, related to Figure 2.**

**A.** PCA plots illustrating variance in RNA-seq expression data between single FGOs of indicated genotypes (*Kdm2b*$^{\Delta CxxC}$, *Kdm2a*$^{KO}$*Kdm2b*$^{\Delta CxxC}$, *Kdm2a*$^{KO}$*Kdm2b*$^{KO}$, *Ring1*$^{KO}$*Rnf2*$^{KO}$ and respective *ctrl* FGOs).

**B.** Total and mapped RNA-seq read counts of individual FGOs indicated in panel a.

**C.** Reproducibility between replicates of H2AK119u1 CUT&RUN data of *ctrl*, *Kdm2a*$^{KO}$*Kdm2b*$^{\Delta CxxC}$ and *Kdm2a*$^{KO}$*Kdm2b*$^{KO}$ FGOs produced in this study. Likewise, between CUT&RUN data in *ctrl* FGOs of this study (replicates pooled) and publicly available data[36]. Depicted are log2 transformed, library normalized counts over all 5kbp genomic tiles.

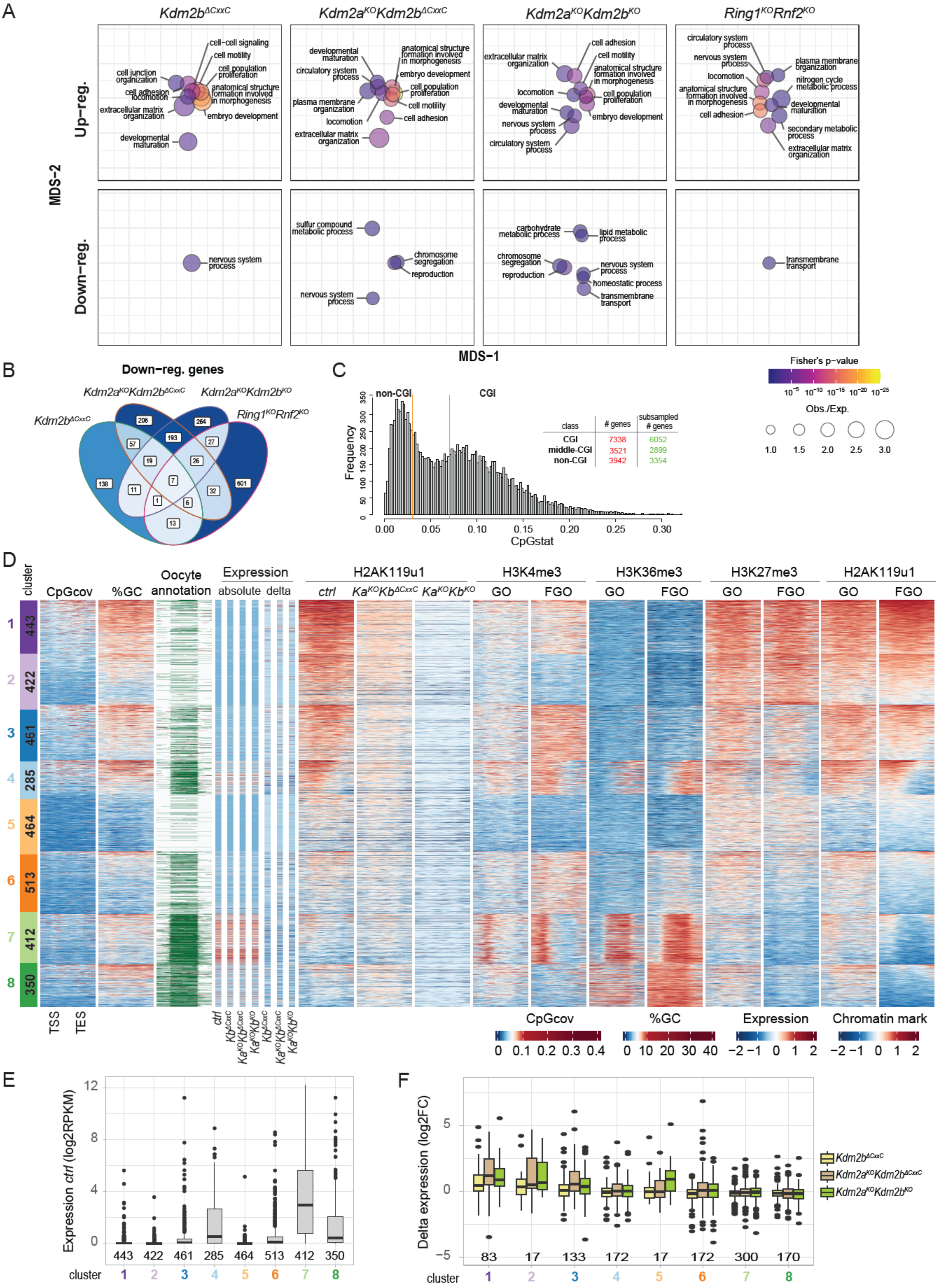

A
Kdm2bΔCxxC
Kdm2aKOKdm2bΔCxxC
Kdm2aKOKdm2bKO
Ring1KORnf2KO
Up−reg.
Down−reg.
MDS−2
MDS−1
cell−cell signaling
cell motility
cell population proliferation
anatomical structure formation involved in morphogenesis
embryo development
cell junction organization
cell adhesion
locomotion
extracellular matrix organization
developmental maturation
circulatory system process
plasma membrane organization
nervous system process
nitrogen cycle metabolic process
secondary metabolic process
sulfur compound metabolic process
chromosome segregation
reproduction
carbohydrate metabolic process
lipid metabolic process
homeostatic process
transmembrane transport
Fisher's p−value
Obs./Exp.
1.0
1.5
2.0
2.5
3.0
B
Down−reg. genes
206
264
193
57
27
19
26
138
7
601
11
32
1
6
13
C
non-CGI
CGI
Frequency
CpGstat
class
### genes
subsampled # genes
CGI 7338 6052
middle-CGI 3521 2899
non-CGI 3942 3354
D
cluster
CpGcov
%GC
Oocyte annotation
Expression
absolute
delta
H2AK119u1
ctrl
H3K4me3
H3K36me3
H3K27me3
GO
FGO
1 443
2 422
3 461
4 285
5 464
6 513
7 412
8 350
TSS
TES
Chromatin mark
E
Expression ctrl (log2RPKM)
443 422 461 285 464 513 412 350
cluster
1 2 3 4 5 6 7 8
F
Delta expression (log2FC)
83 17 133 172 17 172 300 170
cluster
1 2 3 4 5 6 7 8

**Figure S3: KDM2A/KDM2B regulate H2AK119u1 deposition and gene expression during oogenesis, related to Figure 2.**

- **A.** Enrichments of top gene ontology terms for genes up- or down-regulated in *Kdm2a*$^{KO}$*Kdm2b*$^{KO}$, *Kdm2a*$^{KO}$*Kdm2b*$^{\Delta CxxC}$, *Kdm2b*$^{\Delta CxxC}$ ad *Ring1*$^{KO}$*Rnf2*$^{KO}$ FGOs over *ctrl* FGOs. Bubbles representing GO terms are scaled according to enrichments, colored according to statistical significance and positioned relative to one another to reflect similarities between significantly affected genes with corresponding GO terms.
- **B.** Venn diagram showing numbers of genes down-regulated in *Kdm2b*$^{\Delta CxxC}$, *Kdm2a*$^{KO}$*Kdm2b*$^{\Delta CxxC}$, *Kdm2a*$^{KO}$*Kdm2b*$^{KO}$ and/or *Ring1*$^{KO}$*Rnf2*$^{KO}$ FGOs.
- **C.** Distribution of genes according to the CpG density within 500 bp region upstream of their transcriptional start site defined by UCSC. Genes with high and low CpG density are referred to as CGI- and non-CGI promoter genes and have been further sub-selected for those lacking antisense expression during oogenesis. Sub-selected genes have been analyzed in subsequent figures.
- **D.** Heatmap displaying sequence composition, transcriptional and chromatin variables within non-CGI-promoter genes (5kb upstream, TSS, gene body, TES, and 5 kb downstream) grouped into 8 gene clusters by k-means clustering. Clusters 1 to 8 contain 443, 422, 461, 282, 464, 513, 412 and 350 genes. From left to right: CpG coverage; GC percentage; oocyte specific sense (green) and antisense (red) transcripts[40]; absolute RNA (scaled RPKM) in *ctrl*, *Kdm2b*$^{\Delta CxxC}$, *Kdm2a*$^{KO}$*Kdm2b*$^{\Delta CxxC}$ and *Kdm2a*$^{KO}$*Kdm2b*$^{KO}$ FGOs; log2FC expression in *mutant* vs *ctrl* FGOs (delta); H2AK119u1 occupancy in FGOs of indicated genotypes; H3K4me3, H3K36me3, H3K27me3 and H2AK119u1 occupancies in wildtype GOs and FGOs[21,19,36]. All chromatin data are shown as Z-scores. Expression correlates with H3K4me3 promoter occupancy and H3K36me3 gene body occupancy while repression with broad H3K27me3 and H2AK119u1 occupancy in GOs.
- **E.** Boxplot presenting RNA expression levels of non-CGI-promoter genes (in log2RPKM) per gene cluster in *ctrl* FGOs.
- **F.** Boxplot presenting log2FC in expression of non-CGI-promoter genes measured in various *mutant* relative to respective *ctrl* FGOs, indicated per gene cluster.

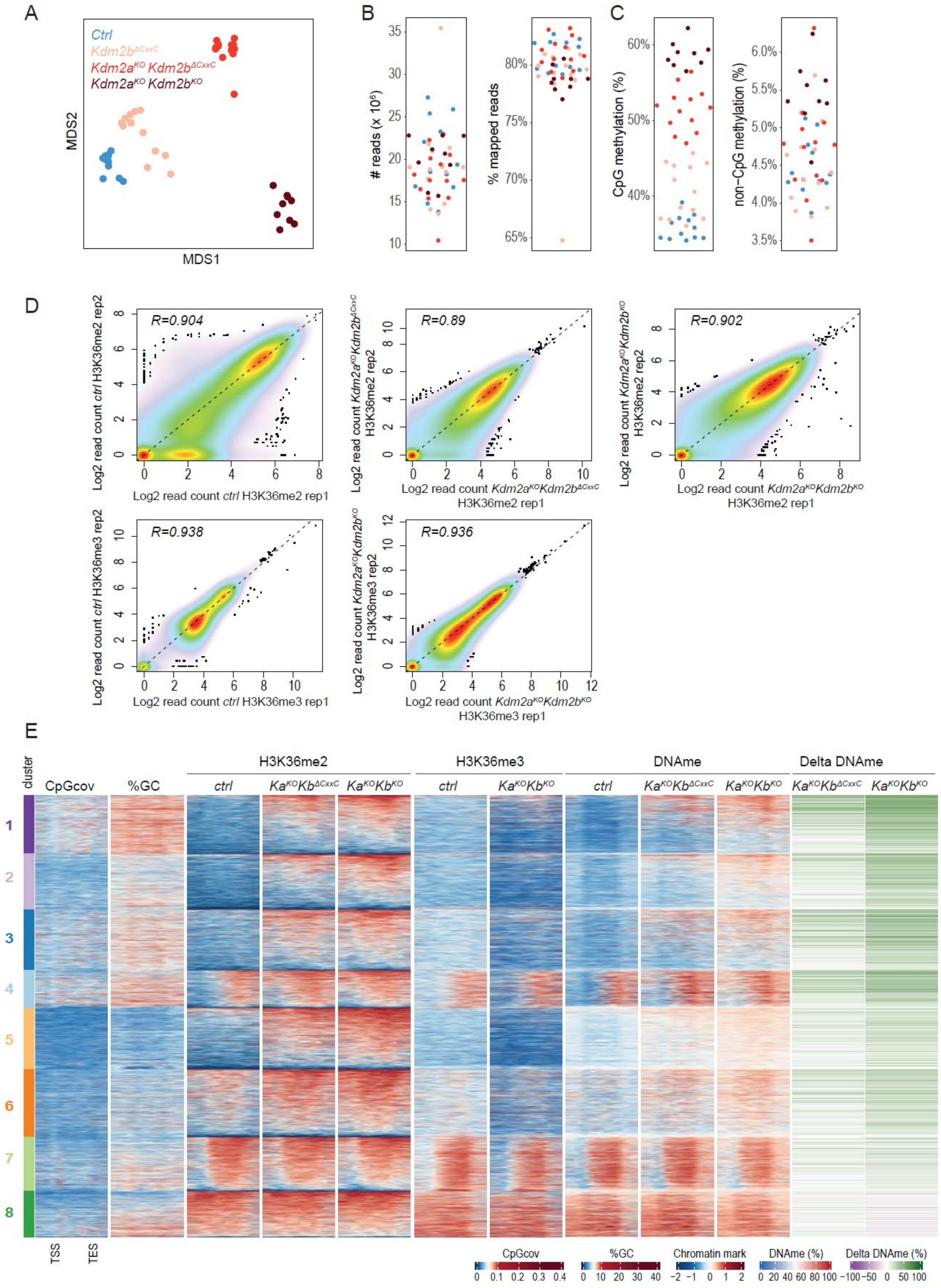
A
Ctrl
Kdm2bΔCxxC
Kdm2aKO Kdm2bΔCxxC
Kdm2aKO Kdm2bKO
MDS2
MDS1
B
### reads (x 10^6)
% mapped reads
C
CpG methylation (%)
non-CpG methylation (%)
D
R=0.904
R=0.89
R=0.902
R=0.938
R=0.936
Log2 read count ctrl H3K36me2 rep1
Log2 read count ctrl H3K36me2 rep2
Log2 read count Kdm2aKOKdm2bΔCxxC H3K36me2 rep1
Log2 read count Kdm2aKOKdm2bΔCxxC H3K36me2 rep2
Log2 read count Kdm2aKOKdm2bKO H3K36me2 rep1
Log2 read count Kdm2aKOKdm2bKO H3K36me2 rep2
Log2 read count ctrl H3K36me3 rep1
Log2 read count ctrl H3K36me3 rep2
Log2 read count Kdm2aKOKdm2bKO H3K36me3 rep1
Log2 read count Kdm2aKOKdm2bKO H3K36me3 rep2
E
cluster
CpGcov
%GC
H3K36me2
H3K36me3
DNAme
Delta DNAme
ctrl
KaKOKbΔCxxC
KaKOKbKO
TSS
TES
Chromatin mark
DNAme (%)
Delta DNAme (%)

**Figure S4: Quality control analyses of WGBS and CUT&RUN-seq data, related to Figure 3.**

- **A.** MDS plot illustrating variance in WGBS data between multiple libraries generated for a given indicated genotype and between genotypes (*Kdm2b*$^{\Delta CxxC}$, *Kdm2a*$^{KO}$*Kdm2b*$^{\Delta CxxC}$, *Kdm2a*$^{KO}$*Kdm2b*$^{KO}$ and *ctrl* FGOs).
- **B.** Total and mapped WGBS-seq read counts of individual libraries indicated in panel **A**.
- **C.** Percentage CpG and non-CpG (CHG, CHH) methylation of libraries indicated in panel **A**.
- **D.** Reproducibility between replicates of H3K36me2 and H3K36me3 CUT&RUN data of *ctrl, Kdm2a*$^{KO}$*Kdm2b*$^{\Delta CxxC}$ and *Kdm2a*$^{KO}$*Kdm2b*$^{KO}$ FGOs produced in this study. Depicted are log2 transformed, library normalized counts over all 5kbp genomic tiles.
- **E.** Heatmap displaying sequence composition and chromatin variables within 8 non-CGI-promoter gene clusters in FGOs, identical to those described in Figure S3D. From left to right: CpG coverage; GC percentage; H3K36me2, H3K36me3 and DNAme in FGOs of indicated genotypes; differential (Delta) DNAme at non-CGI promoters in *Kdm2a*$^{KO}$*Kdm2b*$^{KO}$ and *Kdm2a*$^{KO}$*Kdm2b*$^{\Delta CxxC}$ FGOs.

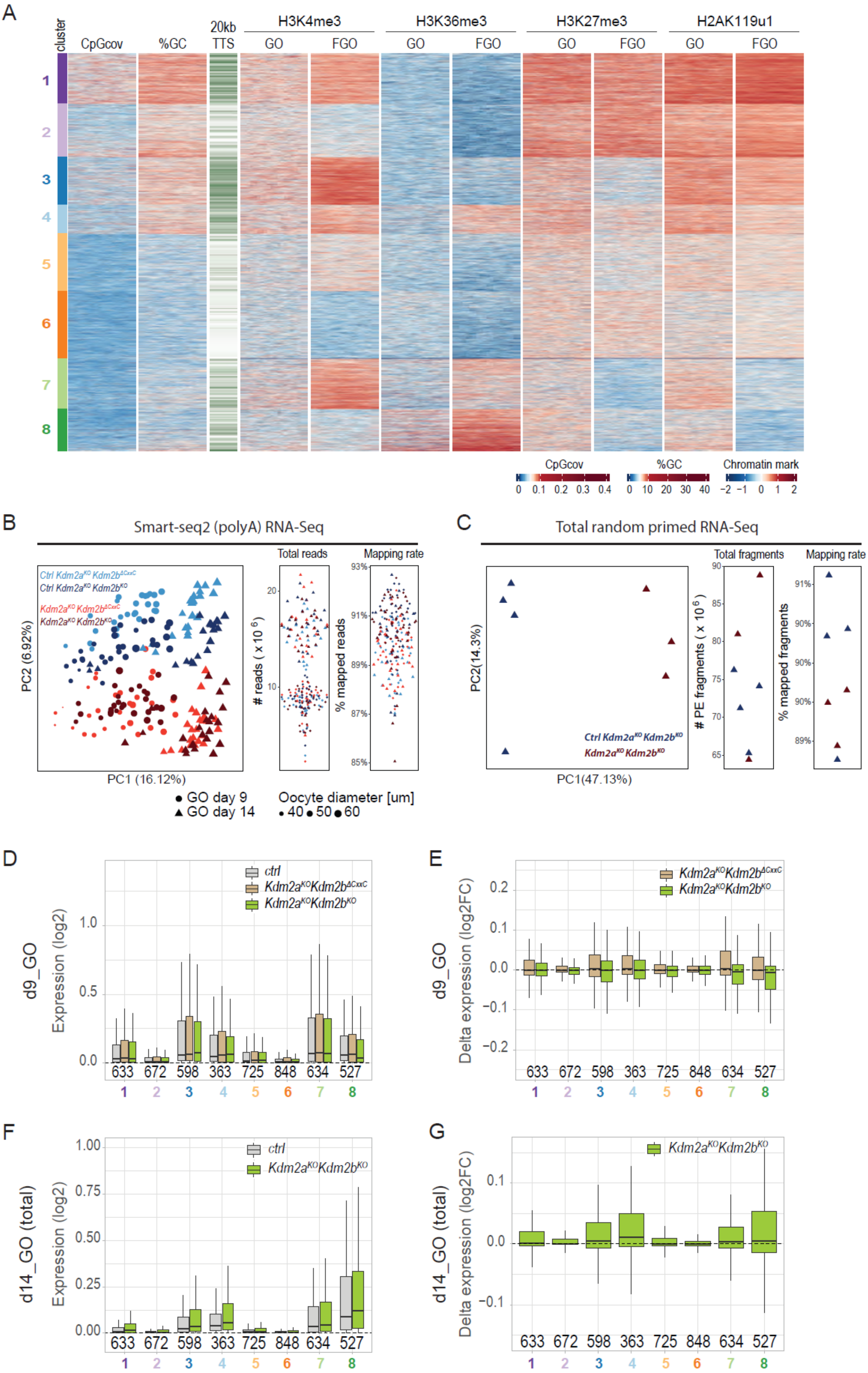
A
B
Smart-seq2 (polyA) RNA-Seq
C
Total random primed RNA-Seq
D
E
F
G

**Figure S5: H3K36me2 and DNAme accumulate throughout the genome of *Kdm2a*/*Kdm2b* mutant oocytes, independently of transcription, related to Figure 4.**

**A.** Heatmap displaying sequence composition and chromatin variables within 8 clusters of 10 kb intergenic regions (20 neighboring 500 bp bins) in oocytes. From left to right: CpG coverage; GC percentage; presence of annotated TTS in 20 kb flanking regions, that could be compatible with run-through transcription through the window; H3K4me3, H3K36me3, H3K27me3 and H2AK119u1 occupancy in wildtype GOs and FGOs[21, 19,36].

**B.** PCA plot illustrating variance in smart-seq2 polyA-based RNA-seq expression data between single GOs at day 9 and 14 of indicated genotypes ($Kdm2a^{KO}Kdm2b^{\Delta CxxC}$, $Kdm2a^{KO}Kdm2b^{KO}$ and respective *ctrl* FGOs). Total and mapped RNA-seq read counts of individual GOs are also indicated.

**C.** PCA plot illustrating variance in total random primed RNA-seq expression data between libraries prepared of pooled $Kdm2a^{KO}Kdm2b^{KO}$ and *ctrl* GOs isolated at day 14 of development. Total and mapped RNA-seq fragment counts per library are indicated as well.

**D.** Boxplot displaying absolute expression for 10 kb intergenic regions in 8 clusters in *ctrl* and $Kdm2a^{KO}Kdm2b^{\Delta CxxC}$ and $Kdm2a^{KO}Kdm2b^{KO}$ GOs at day9. Numbers of regions per cluster are indicated. RNA expression was measured by Smart-seq2 protocol.

**E.** Boxplot displaying log2FC expression for 10 kb intergenic regions in 8 clusters in $Kdm2a^{KO}Kdm2b^{\Delta CxxC}$ and $Kdm2a^{KO}Kdm2b^{KO}$ GOs relative to *ctrl* GOs at day9. Numbers of regions per cluster are indicated. RNA expression was measured by Smart-seq2 protocol.

**F.** Boxplot displaying absolute expression for 10 kb intergenic regions in 8 clusters in *ctrl* and $Kdm2a^{KO}Kdm2b^{KO}$ GOs at day14. Numbers of regions per cluster are indicated. RNA expression was measured by a random-primed based protocol.

**G.** Boxplot displaying log2FC expression for 10 kb intergenic regions in 8 clusters in $Kdm2a^{KO}Kdm2b^{KO}$ GOs relative to *ctrl* GOs at day 14. Numbers of regions per cluster are indicated. RNA expression was measured by a random-primed based protocol.

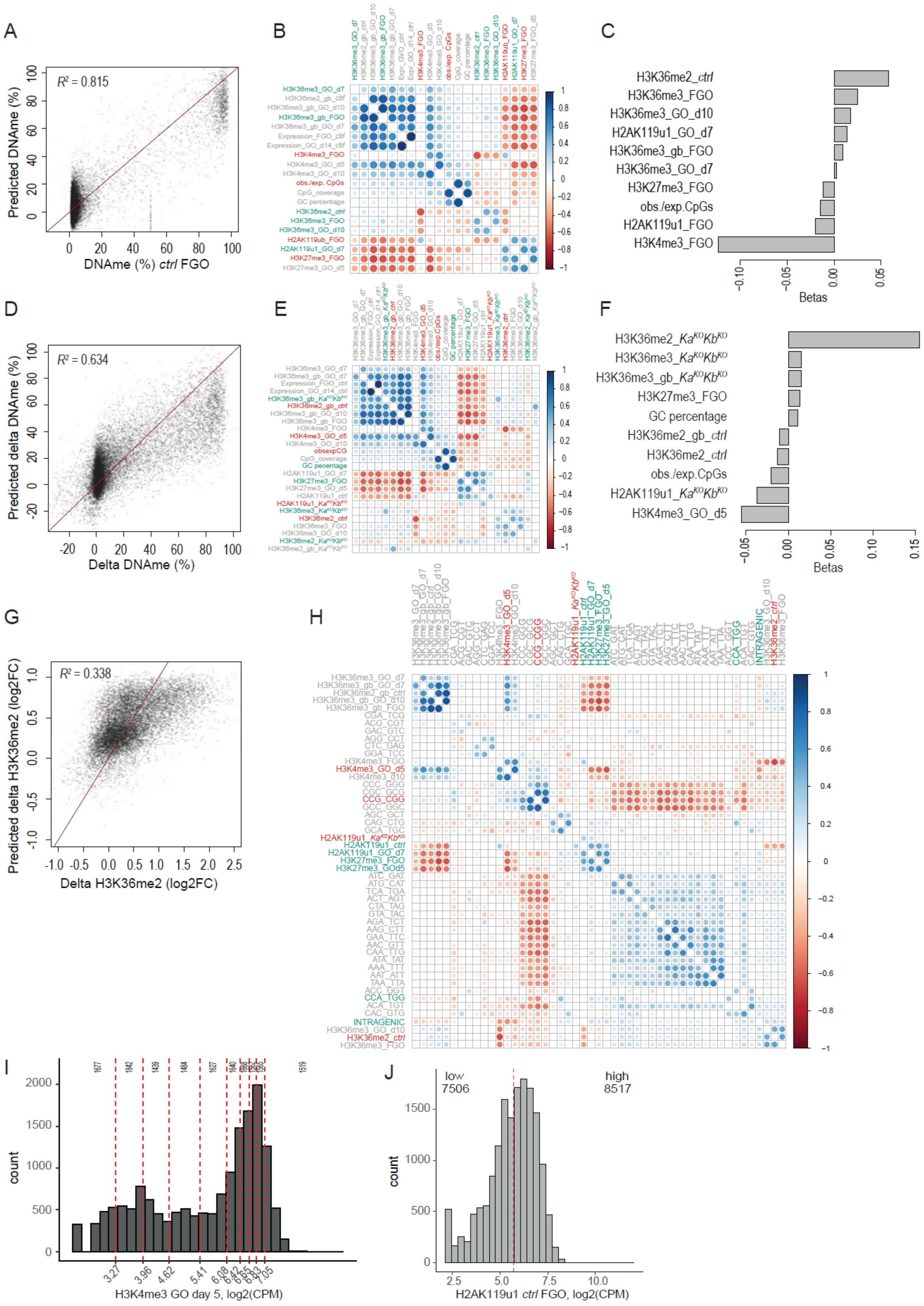
A
B
C
$R^2$ = 0.815
Predicted DNAme (%)
DNAme (%) ctrl FGO
H3K36me2_ctrl
H3K36me3_FGO
H3K36me3_GO_d10
H2AK119u1_GO_d7
H3K36me3_gb_FGO
H3K36me3_GO_d7
H3K27me3_FGO
obs./exp.CpGs
H2AK119u1_FGO
H3K4me3_FGO
Betas
D
E
F
$R^2$ = 0.634
Predicted delta DNAme (%)
Delta DNAme (%)
H3K36me2_KaKOKbKO
H3K36me3_KaKOKbKO
H3K36me3_gb_KaKOKbKO
H3K27me3_FGO
GC percentage
H3K36me2_gb_ctrl
H3K36me2_ctrl
obs./exp.CpGs
H2AK119u1_KaKOKbKO
H3K4me3_GO_d5
Betas
G
H
$R^2$ = 0.338
Predicted delta H3K36me2 (log2FC)
Delta H3K36me2 (log2FC)
I
J
count
H3K4me3 GO day 5, log2(CPM)
low
7506
high
8517
count
H2AK119u1 ctrl FGO, log2(CPM)

**Figure S6: Identifying chromatin and sequence features underlying aberrant H3K36me2 and DNAme acquisition in *Kdm2a*/*Kdm2b* mutant oocytes by regularized linear regression analysis, related to Figure 5.**

**A.** Scatter plot showing DNAme (%) at promoter and intragenic CGIs measured in *ctrl* FGOs versus DNAme (%) at CGIs as predicted as predicted by regularized linear regression analysis using chromatin states in *ctrl* oocytes. $R^2$ = 0.815. Line represents diagonal.

**B.** Dotplot diagram presenting the correlations between sequence, expression and chromatin features at CGIs measured in GOs and FGOs as indicated by the blue to red color gradient. Green and red labeled parameters contribute positively or negatively to the DNAme predictions as described in Figure S6A. Abbreviations: gb = signal in gene body; obs./exp. CpGs = observed over expected CpGs in CGIs; d7, d10: postnatal day 7 or 10.

**C.** Barplot showing the *beta* coefficients of the top 10 variables contributing to the prediction of DNAme status at CGIs in *ctrl* FGOs as described in Figure S6A.

**D.** Scatter plot showing the difference in DNAme (%) at promoter and intragenic CGIs in $Kdm2a^{KO}Kdm2b^{KO}$ over *ctrl* FGOs versus the difference in DNAme (%) between genotypes as predicted by regularized linear regression analysis using chromatin states in *ctrl* and $Kdm2a^{KO}Kdm2b^{KO}$ oocytes. $R^2$ = 0.634. Line represents diagonal.

**E.** Dotplot diagram presenting the correlations between sequence, expression and chromatin features at CGIs measured in GOs and FGOs as indicated by the blue to red color gradient. Green and red labeled parameters contribute positively or negatively to the DNAme predictions as described in Figure S6D. Abbreviations: gb = signal in gene body; obs./expt. CpGs = observed over expected CpGs in CGIs; d5: postnatal day 5.

**F.** Barplot showing the *beta* coefficients of the top 10 variables contributing to the prediction of differential DNAme at CGIs in $Kdm2a^{KO}Kdm2b^{KO}$ over *ctrl* FGOs as described in Figure S6D.

**G.** Scatter plot showing the difference in H3K36me2 occupancy at promoter and intragenic CGIs in $Kdm2a^{KO}Kdm2b^{KO}$ FGOs over *ctrl* FGOs versus the difference in H3K36me2 occupancy between genotypes as predicted by regularized linear regression analysis using chromatin states in *ctrl* and $Kdm2a^{KO}Kdm2b^{KO}$ oocytes and trinucleotide sequences. $R^2$ = 0.338. Line represents diagonal.

**H.** Dotplot diagram presenting the correlations between sequence, expression and chromatin features at CGIs measured in growing and FGOs as indicated by the blue to red color gradient. Green and red labeled parameters contribute positively or negatively to the H3K36me2 predictions as described in Figure S6G. Abbreviations: gb = signal in gene body; obs./expt. CpGs = observed over expected CpGs in CGIs; Trinucleotide frequencies variables are encoded as FFF_RRR pairs where RRR is the reverse complement of FFF.

**I.** Histogram displaying distribution of H3K4me3 occupancy levels at 533 bp-regions surrounding 16’023 CGIs in GOs. Classification of CGIs into 10 bins with approximately equal numbers of CGIs is indicated.

**J.** Histogram displaying distribution of H2AK119u1 occupancy levels at 533 bp-regions surrounding 16’023 CGIs in GOs. CGIs classified as having low or high H2K119u1 levels are indicated.

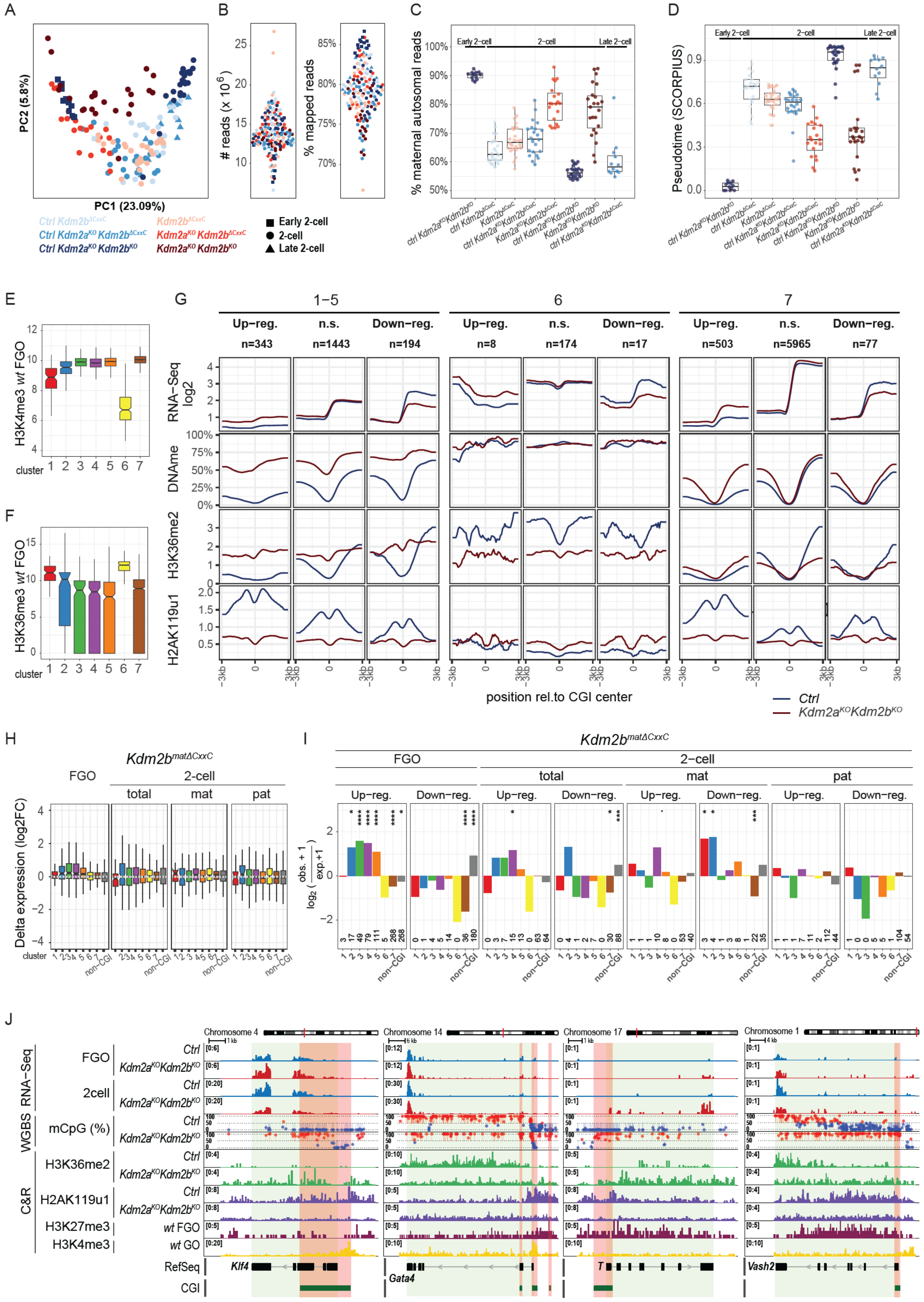

A
PC1 (23.09%)
PC2 (5.8%)
Ctrl Kdm2b^ΔCxxC
Ctrl Kdm2a^KO Kdm2b^ΔCxxC
Ctrl Kdm2a^KO Kdm2b^KO
Kdm2b^ΔCxxC
Kdm2a^KO Kdm2b^ΔCxxC
Kdm2a^KO Kdm2b^KO
B
### reads (x 10^6)
% mapped reads
Early 2-cell
2-cell
Late 2-cell
C
% maternal autosomal reads
D
Pseudotime (SCORPIUS)
E
H3K4me3 wt FGO
cluster
F
H3K36me3 wt FGO
G
Up−reg.
n.s.
Down−reg.
RNA−Seq log2
DNAme
H3K36me2
H2AK119u1
position rel.to CGI center
Ctrl
Kdm2a^KO Kdm2b^KO
H
Kdm2b^matΔCxxC
FGO
2-cell
total
mat
pat
Delta expression (log2FC)
non-CGI
I
log2 (obs. + 1 / exp.+1)
J
RNA-Seq
WGBS
C&R
FGO
2cell
mCpG (%)
H3K36me2
H2AK119u1
H3K27me3
H3K4me3
wt FGO
wt GO
RefSeq
CGI
Klf4
Gata4
T
Vash2
Chromosome 4
Chromosome 14
Chromosome 17
Chromosome 1

**Figure S7: RNA and chromatin dynamics in *Kdm2a*/*Kdm2b* mutant versus *ctrl* FGOs at UCSC-defined CGIs associated with genes being up- or down-regulated in 2-cell embryos, related to Figure 7.**

**A.** PCA plot illustrating variance in RNA-seq expression data between single 2-cell embryos of indicated genotypes (*Kdm2b*$^{\Delta CxxC}$, *Kdm2a*$^{KO}$*Kdm2b*$^{\Delta CxxC}$, *Kdm2a*$^{KO}$*Kdm2b*$^{KO}$, and respective *ctrl*) and stage (early 2-cell, 2-cell and late 2-cell stage).

**B.** RNA-seq read counts and mapping rates for individual 2-cell embryos in Figure S7A.

**C.** Percentage of SNP based allelic reads attributed as transcribed from the maternal genome for single embryos grouped by stage and genotype.

**D.** Inferred pseudotime for single embryos grouped by stage and genotype.

**E.** **F.** Boxplots displaying log2 enrichment for H3K4me3[21] (**E**) and H3K36me3[19] (**F**) in wild-type FGOs at promoter CGIs according to UCSC for genes belonging to the different DNAme clusters, as defined in Figure 7A.

**G.** Metaprofiles for RNA-seq read coverage (including coverage from exon-exon junctions in spliced reads), mCpG methylation, enrichments for H3K36me2 and H2AK119ub1 in *Ctrl* and *Kdm2a*$^{KO}$*Kdm2b*$^{KO}$ FGOs around promoter CGIs belonging to clusters 1-5, 6 and 7 in Figure 7A nearby up-regulated, not significantly changed (n.s.) and down-regulated genes in *Kdm2a*$^{KO}$*Kdm2b*$^{KO}$ FGOs. Direction from negative to positive positions around CGI centers coincide with direction from promoters to gene bodies of nearby genes.

**H.** Boxplots showing log2FC expression of different clusters of CGI-promoter and all non-CGI-promoter genes in *Kdm2b*$^{\Delta CxxC}$ over *ctrl* FGOs and in *Kdm2b*$^{mat\Delta CxxC}$ over *ctrl* 2-cell embryos according to all, mat and pat specific sequencing reads.

**I.** Barplot showing over-/under-representation and statistical significance of CGI-promoter genes belonging to the different DNAme clusters and being either up- or down-regulated in *Kdm2b*$^{\Delta CxxC}$ relative to *ctrl* FGOs and/or in *Kdm2b*$^{mat\Delta CxxC}$ relative to *ctrl* 2-cell embryos for all, mat and pat specific sequencing reads.

**J.** Genomic snapshots of *Klf4*, *Gata4*, *Brachyury* (*T*), and *Vash2* [55] as representative genes gaining DNAme at CGI promoters (highlighted in orange). RNA expression, DNA methylation and chromatin marks are indicated.

**Supplementary Table legends**

**Table S1: Gene expression values in oocytes at d9, d14 and FGO stage for *Kdm2b*$^{\Delta CxxC}$, *Kdm2a*$^{KO}$*Kdm2b*$^{\Delta CxxC}$, *Kdm2a*$^{KO}$*Kdm2b*$^{KO}$, *Ring1*$^{KO}$*Rnf2*$^{KO}$ mutants and respective controls, related to Figures 2, 7, S2, S3 and S7.**

**Table S2: Differential gene expression analysis of oocytes at d9, d14 and FGO stage for *Kdm2b*$^{\Delta CxxC}$, *Kdm2a*$^{KO}$*Kdm2b*$^{\Delta CxxC}$, *Kdm2a*$^{KO}$*Kdm2b*$^{KO}$, *Ring1*$^{KO}$*Rnf2*$^{KO}$ mutants and respective controls, related to Figures 2, 7, S2, S3 and S7.**

**Table S3: DNA methylation levels at CpG islands for *Kdm2b*$^{\Delta CxxC}$, *Kdm2a*$^{KO}$*Kdm2b*$^{\Delta CxxC}$, *Kdm2a*$^{KO}$*Kdm2b*$^{KO}$ and *ctrl* FGOs, related to Figures 7, S4 and S7.**

**Table S4: Expression values for total, maternal and paternal gene transcripts for *Kdm2b*$^{\Delta CxxC}$, *Kdm2a*$^{KO}$*Kdm2b*$^{\Delta CxxC}$, *Kdm2a*$^{KO}$*Kdm2b*$^{KO}$ and respective control 2-cell embryos, related to Figures 6, 7 and S7.**

**Table S5: Differential expression analysis of total, maternal and paternal gene transcripts for *Kdm2b*$^{\Delta CxxC}$, *Kdm2a*$^{KO}$*Kdm2b*$^{\Delta CxxC}$, *Kdm2a*$^{KO}$*Kdm2b*$^{KO}$ and respective control 2-cell embryos, related to Figures 6, 7 and S7.**

**Table S6: GO-terms for CGI-promoter genes belonging to clusters in Figure 7A.**

**Table S7: Gene Ontology terms for selected key developmental functions in early embryos, related to Figure 7.**
